# Removal of repressive histone marks creates epigenetic memory of recurring heat in *Arabidopsis*

**DOI:** 10.1101/2020.05.10.086611

**Authors:** Nobutoshi Yamaguchi, Satoshi Matsubara, Kaori Yoshimizu, Motohide Seki, Kouta Hamada, Mari Kamitani, Yuko Kurita, Soichi Inagaki, Takamasa Suzuki, Eng-Seng Gan, Taiko To, Tetsuji Kakutani, Atsushi J. Nagano, Akiko Satake, Toshiro Ito

## Abstract

Acclimation to high temperature increases plants’ tolerance of subsequent lethal high temperatures^1-3^. Although epigenetic regulation of plant gene expression is well studied, how plants maintain a memory of environmental changes over time remains unclear. Here, we show that JUMONJI (JMJ) proteins^4-8^, demethylases involved in histone H3 lysine 27 trimethylation (H3K27me3), are necessary for *Arabidopsis thaliana* heat acclimation. Acclimation induces sustained H3K27me3 demethylation at key *HEAT SHOCK PROTEIN* (*HSP*) loci by JMJs, poising the *HSP* genes for subsequent activation. Upon sensing heat after a 3-day interval, JMJs directly reactivate *HSP* genes. Finally, *jmj* mutants fail to maintain heat memory under fluctuating field temperature conditions. Our findings of an epigenetic memory mechanism involving histone demethylases may have implications for environmental adaptation of field plants.

## Main

The ability to adapt to environmental changes is essential for plants, as sessile organisms, to survive^1^. Plants memorize heat experience over several days and develop future responsiveness^2^. Exposure to moderate temperature enables plants to acquire thermotolerance for subsequent lethal high temperature^3^. In *Arabidopsis thaliana*, HEAT SHOCK TRANSCRIPTION FACTOR A2 (HSFA2) is necessary for the maintenance of acquired thermotolerance^9, 10^. *HEAT SHOCK PROTEIN* (*HSP*) genes encode molecular chaperones that protect cellular proteins from denaturation. Upon sensing of high temperature, the transient binding of HSFA2 at *HSPs* governs the sustained increase of histone marks such as histone H3 lysine 4 trimethylation (H3K4me3) and the expression of the *HSP* genes^11-13^. Then, the expression of heat-memory genes declines gradually while H3K4me3 levels remain high. In that situation, proper maintenance of repressive histone marks should also play important roles in the downregulation of *HSP* genes^13^. Despite the importance of histone modification enzymes, little is known about the underlying mechanism of those enzymes in flexible and reversible *HSP* gene expression.

JUMONJI (JMJ) proteins^4-8^, demethylases involved in histone H3 lysine 27 trimethylation (H3K27me3), are evolutionarily conserved and regulate diverse biological processes. In this study, we investigate the role these H3K27me3 demethylases on acquired thermotolerance in response to recurring heat. We demonstrate that JMJ proteins keep repressive histone marks at low levels on chaperone-encoding small *HSP*s^11-13^ that function as memory genes. Using inducible JMJs and mutants of small *HSP*s, we demonstrate the underlying cause of heat memory is the lower H3K27me3 on small *HSPs*. This histone modification-based transcriptional memory is well-aligned with a mathematical stochastic model that we developed, which predicts expression levels of *sHSP*s. Moreover, we recapitulate fluctuating temperature conditions and suggest that JMJ-mediated sustained H3K27me3 demethylation on small *HSPs* controls heat memory under the conditions.

## Results

### Heat memory defects in H3K27me3 demethylase mutants

To gain insight into the role of histone modification enzymes in the maintenance of heat memory, we focused on a group of H3K27me3 demethylases, the Jumonji-C-domain-containing proteins. Among the 21 JUMONJI (JMJ) proteins, five reportedly possess H3K27me3 demethylation activity: JMJ30, JMJ32, JMJ11/EARLY FLOWERING 6 (ELF6), JMJ12/RELATIVE OF EARLY FLOWERING (REF6), and JMJ13^4-8^. The *ref6-3 elf6C jmj13G* triple mutants exhibit multiple morphological differences from the wild type^8^, including dwarfism, which could have a secondary effect on heat responses; importantly, we observed no differences in leaf size between the 10-day-old wild-type and *jmj30-2 jmj32-1* double and *jmj30-2 jmj32-1 elf6-1 ref6-3* quadruple mutant (hereafter *jmjd* and *jmjq*, respectively) seedlings (Extended Data Fig. 1). Therefore, we examined the role of H3K27me3 demethylases in heat memory using these single and multiple mutants (Fig. 1a–i). When grown at 22°C (the control condition), the wild-type and all mutant plants including *jmjq* survived normally (Fig. 1a–c,h,i). Exposure to heat shock at 44°C (+HS) led to significant reduction of survival rates in both groups of plants (Fig. 1d,e,h,i). Approximately 80% of acclimatized seedlings of the wild type and all mutants except *jmjq* survived (Fig. 1f–i), and they had similar chlorophyll contents and fresh weights (Fig. 1f–i, and Extended Data Fig. 2). By contrast, *jmjq* mutants showed significantly lower survival rate, chlorophyll content, and fresh weights after acclimation and heat shock (+ACC +HS) than the wild type or any other mutant combination (Fig. 1f–i, and Extended Data Fig. 2). Thus, JMJ demethylases are redundantly required for heat acclimation.

**Figure 1.**
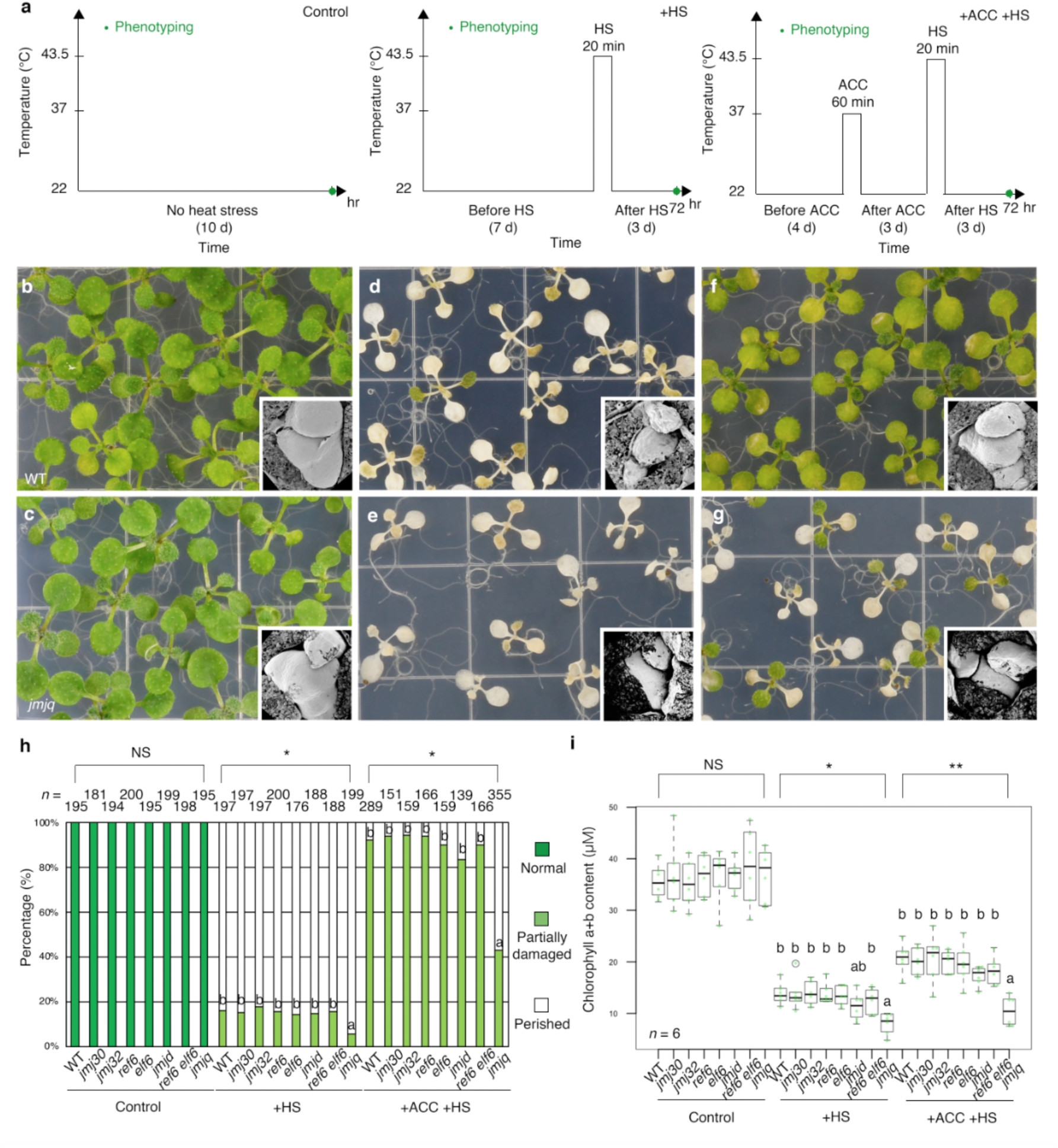
H3K27me3 demethylase activity is required for acquired thermotolerance. **a**, Schematic representation of the temperature conditions used. Phenotype, green. Left, normal plant growth condition. Center, basal thermotolerance condition. Right, heat-stress memory condition. **b** and **c**, Wild type (b) and *jmj30 jmj32 ref6 elf6* quadruple mutants (*jmjq*) (c) grown under control conditions. **d** and **e**, Wild-type (d) and *jmjq* (e) grown under +HS conditions. **f** and **g**, Wild type (f) and *jmjq* (g) grown under +ACC +HS conditions. Insets show scanning electron microscope images of representative shoot apical meristems. **h**, Quantification of survival rate. 10-day-old seedlings grown under the three different temperature conditions were categorized into three groups based on phenotypic severity: green, normal growth; light green, partially damaged; white, perished. Significance was determined by χ^2^ test followed by post-hoc test. *n* > 138. **i**, Quantification of chlorophyll contents. Light green jitter dots and white circles represent the chlorophyll contents from each sample and from statistical outliers, respectively. One-way ANOVA test, \**p* < 1.0 × 10^−3^, \*\**p* < 1.0 × 10^−4^. Different letters indicate significant differences, while the same letters indicate non-significant differences based on post-hoc Tukey’s HSD test. *p* < 0.05. NS, nonsignificant. *n* = 6.

**Figure 2.**
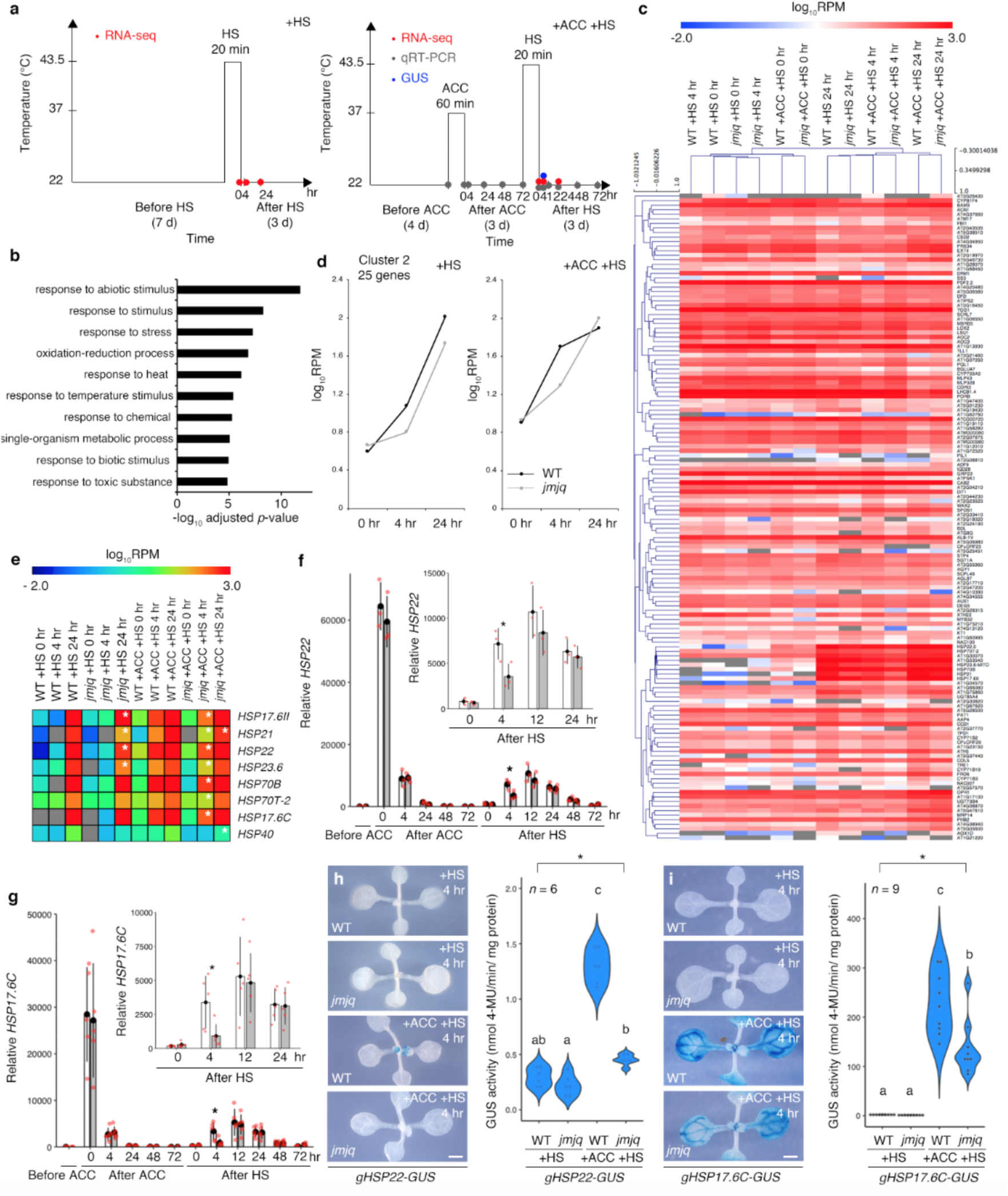
Histone H3K27me3 demethylases mediate rapid reactivation of *HSP* genes during heat acclimation. **a**, Schematic representation of the temperature conditions used. Left, basal thermotolerance condition; right, heat-stress memory condition. RNA-seq, red; qRT-PCR, gray; GUS staining, blue. **b**, GO term enrichment analysis of 142 differentially expressed genes. The top 10 terms determined by their –log_10_-adjusted *p*-values are shown. **c**, A *k*-means clustering of genes differentially expressed between acclimated wild type and *jmjq* mutants 0, 4 or 24 hrs after a tester heat stress. The FDR was <0.05. **d**, Gene expression over time in cluster 2 under basal thermotolerance (left) and heat-stress memory (right) conditions. **e**, Heatmap of the *HSP* genes in cluster 2. White asterisks indicate significant differences. FDR < 0.05. **f** and **g**, qRT-PCR verification of the *HSP22* (f) and *HSP17*.*6C* (g) levels in wild type and *jmjq* mutants grown under the conditions shown in Fig. 2a. Red jitter dots represent expression level from each sample. Asterisks indicate significant difference at 0.05 levels based on Student’s *t*-test between wild type and *jmjq* mutants at the same time pont. **h** and **i**, *gHSP22-GUS* (h) and *gHSP17*.*6C-GUS* (i) expression in wild type and *jmjq* mutants grown under conditions shown in Fig. 2a. Left, Accumulation of beta-glucuronidase. Right, Quantification of GUS activity by MUG assay. One-way ANOVA test, \**p* < 0.05. Different letters indicate significant differences, while the same letters indicate non-significant differences based on post-hoc Tukey’s HSD test (*p* < 0.05). NS, nonsignificant. *n* > 5.

### H3K27me3 demethylases prime activation of heat memory genes

Next, we conducted RNA sequencing (RNA-seq) to identify JMJ-dependent targets whose expression changed in response to acclimation and heat shock (Fig. 2a, and Extended Data Fig. 3). Acclimatized plants often respond to the triggering stress faster, earlier, and/or more strongly than non-acclimatized plants^2^. We screened for genes that showed no difference in expression just after HS in acclimatized wild-type and *jmjq* mutant seedlings, but showed significant differences by either 4 or 24 hrs after HS. These analyses identified 142 genes that were differentially expressed in the *jmjq* mutant as compared to the wild type (FDR < 0.05): 62 upregulated and 80 downregulated in *jmjq* (Supplementary Table 1,2). As expected, stress-related Gene Ontology (GO) terms such as “response to stress” and “response to heat” were significantly enriched among these genes (Fig. 2b and Supplementary Table 3). Clustering analysis using wild-type and *jmjq* data revealed six clusters based on expression patterns (Fig. 2c,d, and Supplementary Table 4). Among the six clusters, cluster 2 (25 genes) showed rapidly increasing expression over the time course after HS in acclimatized wild-type seedlings (Fig. 2d and Extended Data Fig. 4), but this rapid induction was compromised in acclimatized *jmjq* mutants (Fig. 2d). This cluster included eight differentially expressed *HSP* genes whose products function as molecular chaperones (Fig. 2d,e, and Supplementary Table 2). Among them, small *HSP* genes are known for their importance in heat acclimation, such as *HSP22* and *HSP21*^14, 15^. In the acclimatized wild-type plants, *HSP22, HSP17*.*6C*, and *HSP21* were immediately induced at high levels upon ACC and moderately activated within 4 hrs after HS (Fig. 2f,g, and Extended Data Fig. 5). This moderate and delayed activation of *HSP* genes upon HS may correspond to recovery time due to a reduced cell viability following the HS^16^ (Extended Data Fig. 6). The expression of those three *HSP* genes at 4 hrs after HS was significantly lower in acclimatized *jmjq* mutants than in acclimatized wild-type seedlings (Fig. 2f,g, and Extended Data Fig. 5). Furthermore, the accumulation of HSP22, HSP17.6C, and HSP21 proteins was lower in acclimatized *jmjq* mutants compared to the wild type 4 hrs after heat shock^17, 18^ (Fig. 2h and i, and Extended Data Fig. 7). These data indicate that JMJ plays an important role in the rapid induction of small *HSP* genes.

**Figure 3.**
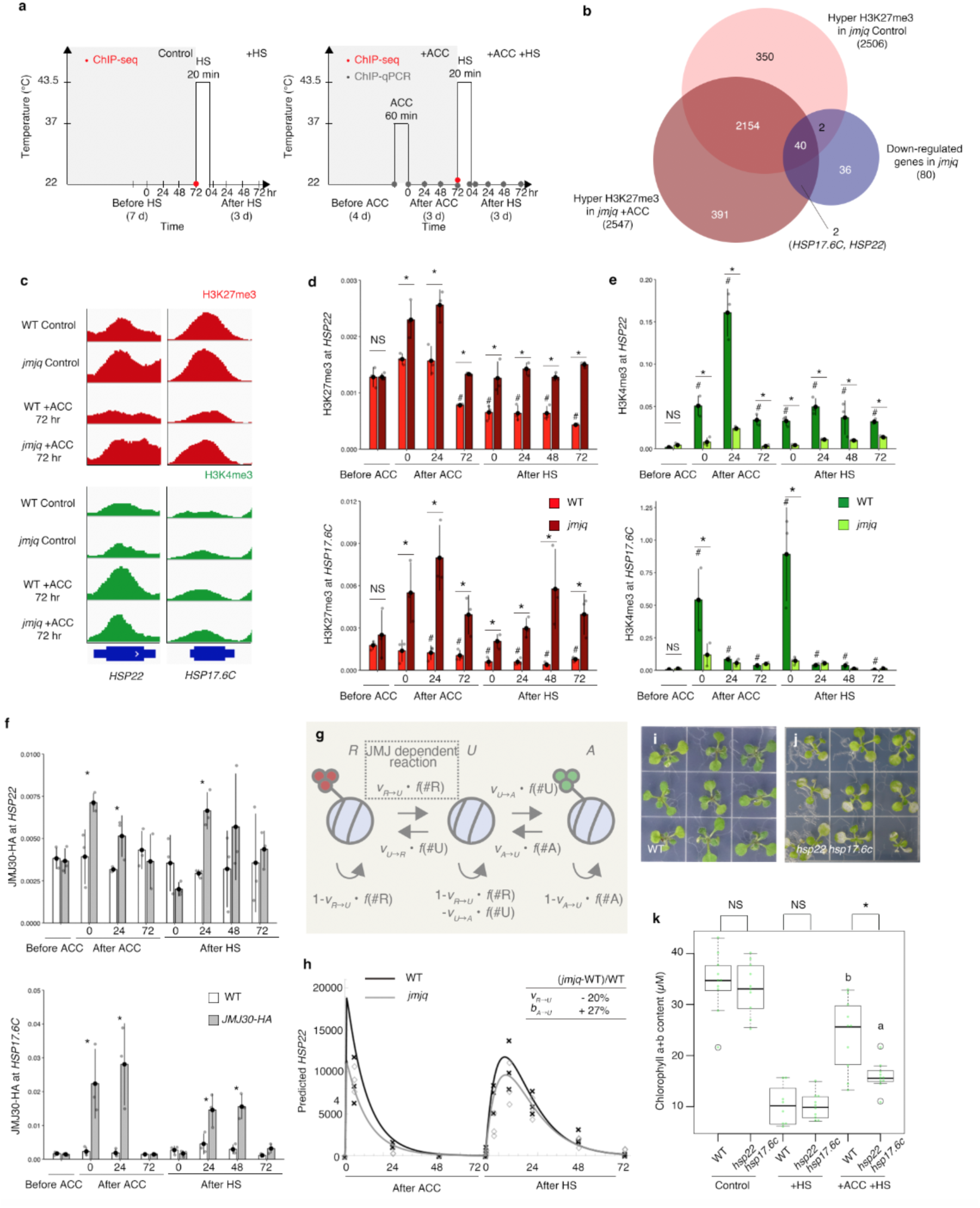
JUMONJI demethylases remove histone H3K27me3 from key heat-memory genes in response to heat. **a**, Schematic representation of the temperature conditions used. ChIP-seq, red; ChIP-qPCR, gray. Left, basal thermotolerance condition; right, heat-stress memory condition. **b**, Venn diagrams showing the overlap between genes downregulated in *jmjq* mutants and genes with elevated H3K27me3 in the mutant with and without acclimation (*p* = 6.2 × 10^−26^ for elevated H3K27me3). **c**, H3K27me3 peaks at the *HSP* genes in wild type and *jmjq* mutants without and with acclimation. **d**-**f**, H3K27me3 (d) and H3K4me3 (e) levels in wild type and *jmjq* mutants, and JMJ30-HA (f) levels, as determined by ChIP-qPCR. Gray jitter dots represent expression level from each sample. Student’s *t*-test compared to the wild type before ACC, ^#^*p* < 0.05. Student’s *t*-test between wild type and *jmjq* mutants at the same time pont, \**p* < 0.05. NS, not significant. **g**, Diagram of the mathematical model capturing the histone modification process. **h**, Profile of *HSP22* expression in wild type and *jmjq* mutants predicted by the model under the conditions shown in Fig. 3a. Experimental data from wild type (crosses) and *jmjq* mutants (diamonds) are also plotted. **i** and **j**, Wild type (**i**) and *hsp22 hsp17*.*6c* mutants (**j**) grown under +ACC +HS condition. **k**, Quantification of chlorophyll contents. One-way ANOVA test, \**p* < 0.05. Different letters indicate significant differences, while the same letters indicate non-significant differences based on post-hoc Tukey’s HSD test (*p* < 0.05). NS, nonsignificant. *n* > 7.

**Figure 4.**
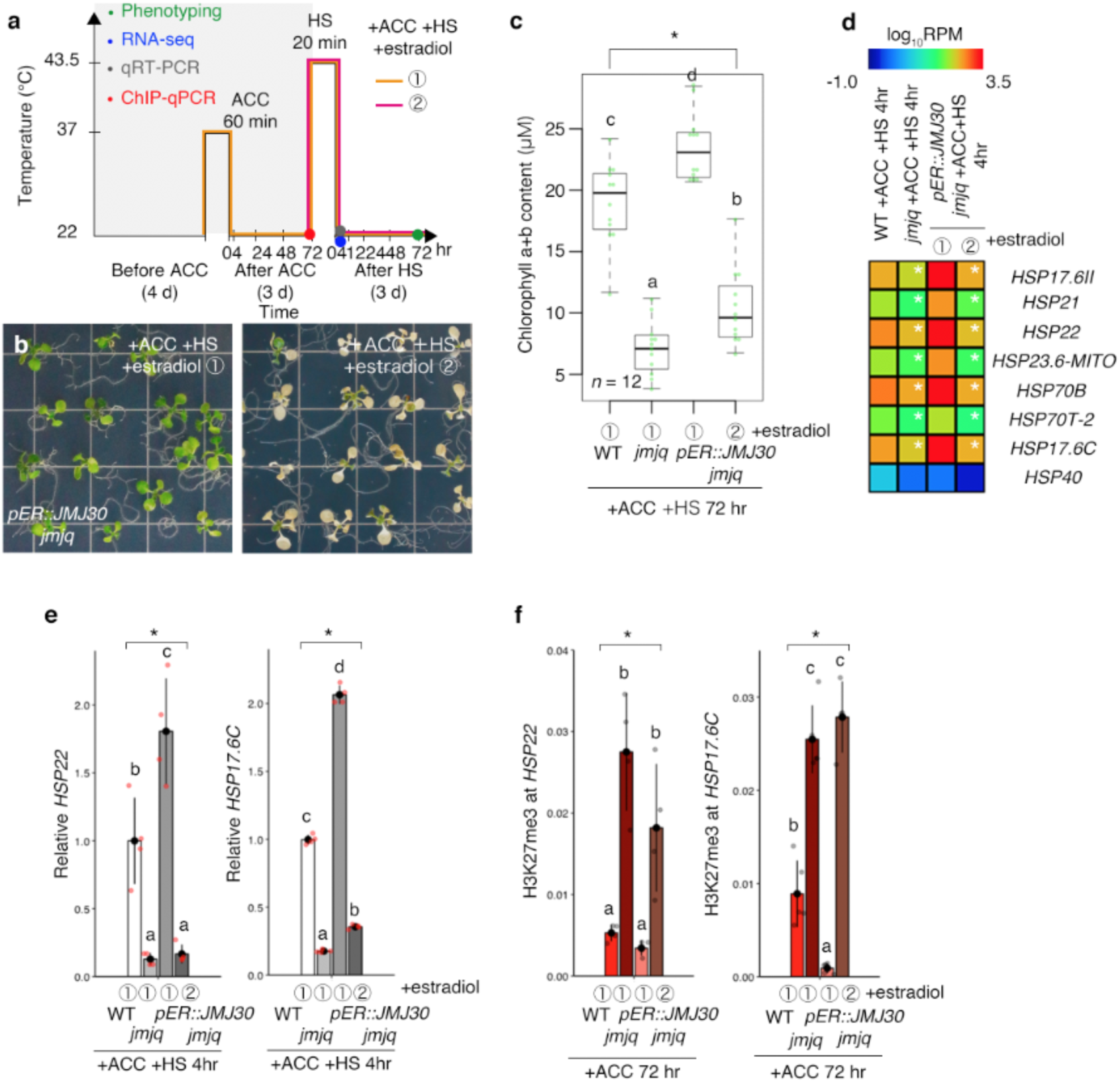
JUMONJI demethylases are required for the removal of histone H3K27me3 from heat-memory genes before acclimation. **a**, Schematic representation of heat-stress memory conditions. Phenotyping, green; RNA-seq, blue; qRT-PCR, gray; ChIP-qPCR, blue. Orange and magenta lines show two different timing of β-estradiol application. **b**, *pER8::JMJ30* transgenic plants in the *jmjq* mutant background subjected to β-estradiol application before acclimation (➀, left) and before heat shock (➁, right). Plants were grown under heat-stress memory condition. **c**, Quantification of chlorophyll contents in plants shown in Fig. 4b.One-way ANOVA test, \**p* < 0.05. Letters above bars indicate significant difference, while the same letters indicate non-significant differences on post-hoc Turkey HSD test (*p* < 0.05). *n* = 12. **d**, Heatmap of the *HSP* genes downregulated in *jmjq* mutants. White asterisks indicate significant differences. The FDR was <0.05. **e** and **f**, Gene expression levels (e) and H3K27me3 levels (f) of *HSP22* and *HSP17*.*6C* in wild type, *jmjq* mutants, and *ER8::JMJ30* transgenic plants in the *jmjq* mutant background subjected to β-estradiol application before acclimation (➀) and before heat shock (➁). One-way ANOVA test, \**p* < 0.05. Letters above bars indicate significant difference on post-hoc Turkey HSD test (*p* < 0.05).

**Figure 5.**
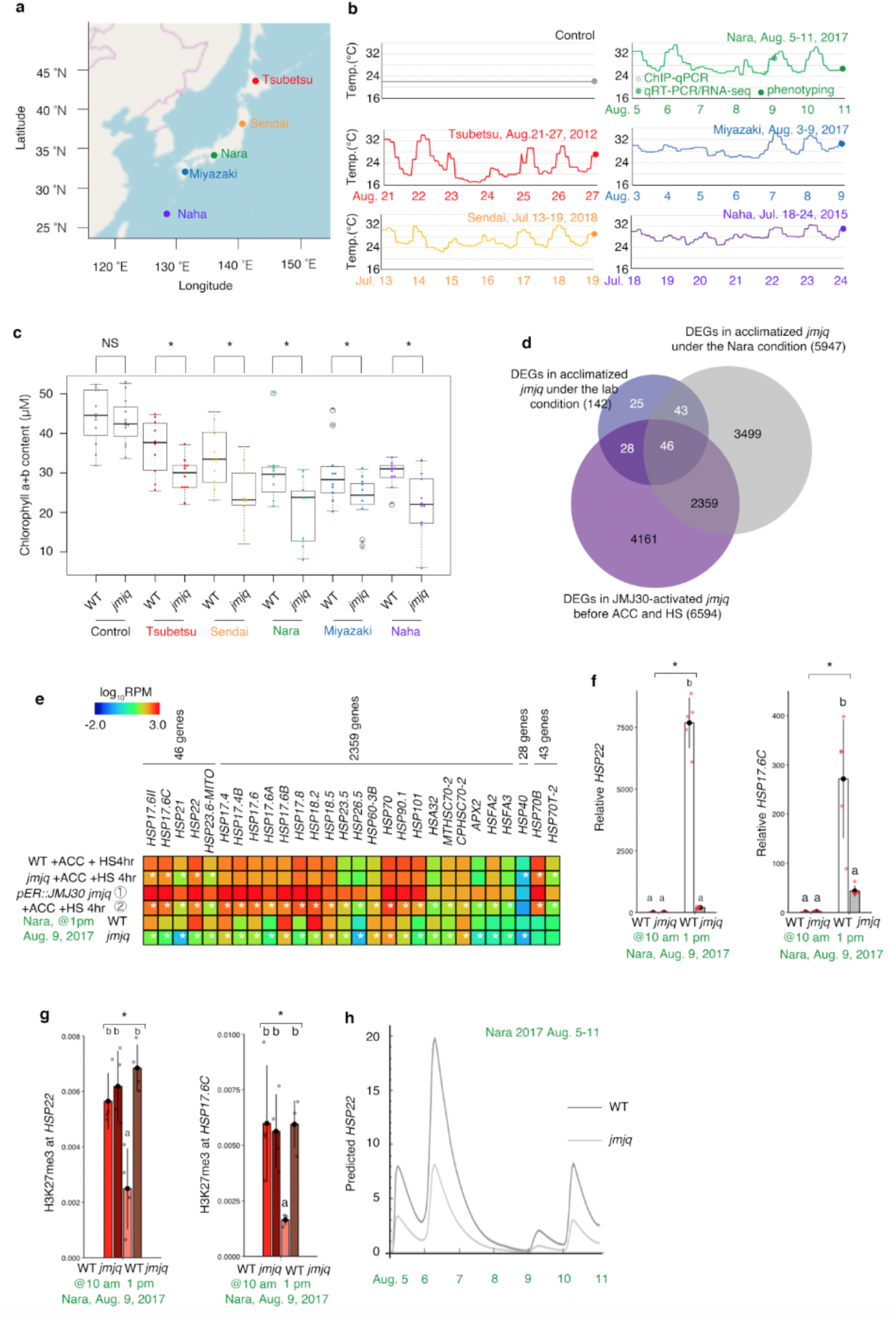
JUMONJI demethylases are required for the removal of histone H3K27me3 from heat-memory genes under fluctuating temperature conditions. **a**, Locations of cities in Japan whose temperature conditions were mimicked to provide a set of naturally varying growth conditions. Red, orange, green, blue, and purple dots represent Tsubetsu, Sendai, Nara, Miyazaki, and Naha, respectively. **b**, Schematic representation of temperature conditions. **c**, Quantification of chlorophyll contents of plants grown under the five conditions shown in Fig. 5b. Colored jitter dots and white circles represent the chlorophyll contents from each sample and statistical outliers, respectively. Student’s *t*-test, \**p* < 0.05). **d**, Venn diagram showing the overlap between genes downregulated in *jmjq* under the lab and Nara conditions. **e**, Heatmap of genes downregulated in *jmjq* mutants under the Nara conditions. White asterisks indicate significant differences. The FDR was <0.05. **f** and **g**, Gene expression levels (f) and H3K27me3 levels (g) of *HSP22* and *HSP17*.*6C* in wild type and *jmjq* mutants grown under the Nara conditions. One-way ANOVA test, \**p* < 0.05. Letters above bars indicate significant difference, while the same letters indicate non-significant differences on post-hoc Turkey HSD test (*p* < 0.05). **h**, Mathematical model prediction for *HSP22* expression under the Nara conditions.

**Figure 6.**
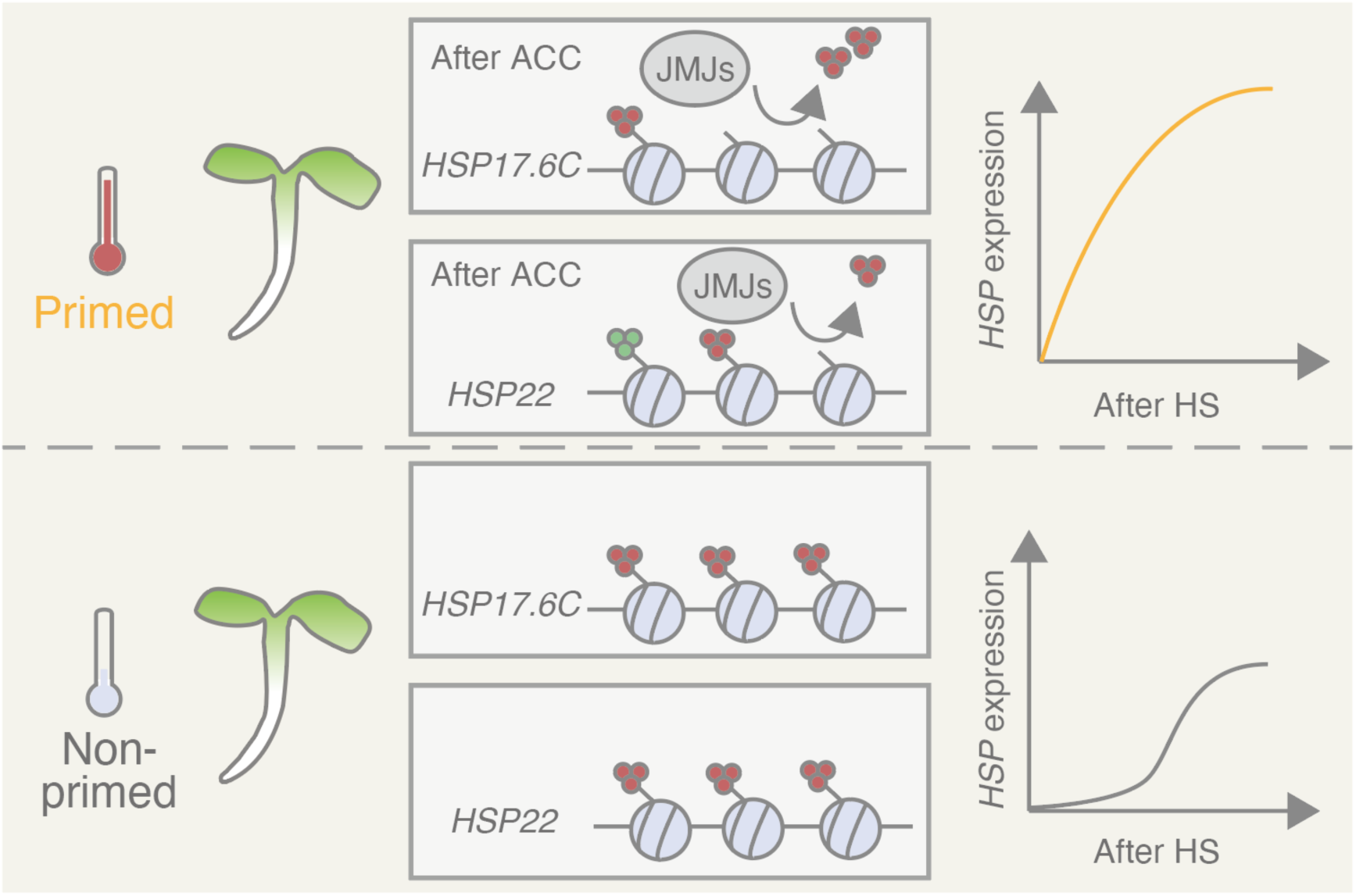
Current model of JUMONJI-mediated heat acclimation. Upon exposure to a triggering heat, plants are acclimatized. JMJ proteins keep H3K27me3 at lower levels on *HSP17*.*6C* and *HSP22*. The primed plants respond to recurring heat faster or stronger than non-primed plants do.

### Sustained H3K27me3 demethylation and H3K4me3 methylation governs transcriptional heat memory

We hypothesized that acclimation-dependent rapid *HSP* activation by JMJ was associated with sustained demethylation of H3K27me3. Based on the phenotypic analysis indicating that the acclimatized wild-type plants memorized heat exposure for at least 3 days, we obtained samples from both wild-type and *jmjq* mutant plants after 3 days of ACC and from equivalent non-acclimatized controls and analyzed them by chromatin immunoprecipitation followed by sequencing (ChIP-seq; Fig. 3a–c). In agreement with a role for JMJ in H3K27me3 removal, more than 2500 genes were gene-body hypertrimethylated by H3K27me3 in *jmjq* compared to the wild type, regardless of whether the conditions compared were non-acclimatized or acclimatized (Fig. 3b, and Extended Data Fig. 8–10). For each condition, 55% of genes identified by RNA-seq as downregulated in *jmjq* also had hypermethylated H3K27me3 (Fig. 3b and Supplementary Table 4)—an overlap significantly greater than that expected by chance (*p* = 6.2 × 10^−26^). Among eight *HSP* genes downregulated in *jmjq* mutants, the *HSP22* and *HSP17*.*6C* loci showed acclimation-dependent removal of H3K27me3 by JMJs (Fig. 3b,c, Extended Data Fig. 11, and Supplementary Table 4,5). ChIP coupled to detection by quantitative polymerase chain reaction (qPCR) confirmed that in the wild type, H3K27me3 levels at the *HSP22* and *HSP17*.*6C* loci were lower following ACC and HS (Fig. 3d,e (#)); in the *jmjq* mutant, however, the levels were significantly higher due to failure of H3K27me3 maintenance (Fig. 3d,e (*) and Extended Data Fig. 11). H3K4me3 levels at *HSP22* and *HSP17*.*6C* increased after ACC and HS in both wild type and *jmjq* mutants (Fig. 3e, Extended Data Fig 11, and Supplementary Table 5,6). Consistent with the levels of H3K27me3 and gene expression of *HSP22*, and *HSP17*.*6C*, JMJ30 directly bound to those genes in response to ACC and HS (Fig. 3f).

To formalize the hidden dynamics of histone modification, we established a mathematical model^19, 20^ (Fig. 3g). Recovery from HS was considered to explain gradual *HSP22* upregulation after HS in both acclimatized wild type and *jmjq* mutants (Fig. 2f and Extended Data Fig. 6). Based on our calculations from the model, *HSP22* expression would be immediately and gradually induced upon ACC and after HS, respectively (Fig. 3h). Furthermore, predicted *HSP22* expression after HS was lower in acclimatized *jmjq* mutants compared to acclimatized wild type (Fig. 3h). Consistent with the biochemical function of JMJs, comparison of the best fit parameter between wild type and *jmjq* mutants indicated that the *jmjq* mutant had a lower transition rate from the repressively modified to the unmodified state (*v*_R→U_), as expected (Fig. 3h, and Supplementary Note). The model highlighted the importance of having multiple modified nucleosomes to lengthen the duration of the memory effect (Extended Data Fig. 12).

In addition, genetic analysis revealed that higher order *hsp22 hsp17*.*6c* double mutants exhibited reduced heat acclimation capacity in terms of chlorophyll content (Fig. 3i–k, and Extended Data Fig. 13). Furthermore, ectopic expression of *HSP17*.*6C* in the *jmjq* mutant led to a partial rescue of the heat-acclimation defects of *jmjq* (Extended Data Fig. 14), similar to the heat tolerance of the *HSP22* overexpressor^15^. Hence, JMJ-mediated demethylation of H3K27me3 at key heat-memory genes acts to control heat acclimation in response to ACC.

### JMJ30 induction creates sustained H3K27me3 demethylation and heat memory

To understand the timing of demethylation activity necessary for heat acclimation, we then introduced a *JMJ30* transgene with an estradiol-inducible promoter into the *jmjq* mutant background^21^ (Fig. 4a). Induction of *JMJ30* before ACC (➀) rescued the *jmjq* mutant phenotype, whereas induction before HS (➁) did not (Fig. 4b,c, and Extended Data Fig. 15). Heat-acclimation-related genes, including seven of the eight *HSP* genes downregulated in *jmjq* mutants, were differentially expressed 4 hrs after HS when *JMJ30* was induced before ACC (Fig. 4d,e, Extended Data Fig. 16,17, and Supplementary Table 7,8). Consistent with that result, after JMJ30 induction in *jmjq* mutants prior to ACC, levels of H3K27me3 at *HSP22* and *HSP17*.*6C* were lower than in the wild type 72 hrs after ACC, while H3K4me3 nor H3 levels stayed unchanged (Fig. 4f and Extended Data Fig. 18,19). Taken together, these results imply that JMJ30 activity is necessary for heat memory prior to acclimation.

### JMJ-mediated sustained H3K27me3 demethylation controls heat memory under fluctuating field conditions

To further examine the key function of JMJs in the adaptation to natural conditions, we grew the wild type and the *jmjq* mutant under fluctuating temperature conditions based on the actual meteorological data from five Japanese cities at different latitudes^22^ (Fig.5a,b, and Supplementary Table 9). Four-day-old seedlings grown under the lab condition of 22°C were transferred to these fluctuating temperature conditions, which included two temperature points >30°C with a 2-day gap. When grown under these five different conditions, the *jmjq* mutant showed significantly lower chlorophyll content than the wild type (Fig. 5c). *HSP17*.*6C, HSP17*.*6II, HSP21, HSP22*, and *HSP23*.*6-MITO* transcripts were reduced in acclimatized *jmjq* mutants grown both under the standard lab condition and under the condition mimicking temperatures in the city of Nara, whereas all five genes were significantly induced when *JMJ30* expression was induced before ACC (Fig.5d,e, Extended Data Fig. 20, and Supplementary Table 10–12). Indeed, many small *HSPs* were differentially expressed in acclimatized *jmjq* mutants under the Nara condition (Fig. 5e). qRT-PCR confirmed that *jmjq* mutants had lower expression of the *HSP22, HSP17*.*6C*, and *HSP21* genes 4 hrs after their exposure to the second heat stress under the Nara condition (Fig. 5f and Extended Data Fig. 21). Furthermore, before the second heat stress, the *jmjq* mutant had higher H3K27me3 at *HSP17*.*6C* and *HSP22* than the wild type (Fig. 5g and Extended Data Fig. 22,23). The stochastic model also predicted the reduction of *HSP* gene expression in acclimatized *jmjq* mutants under the Nara condition (Fig. 5h). Thus, the epigenetic memory mechanism executed by histone demethylases may facilitate environmental adaptation in natural conditions (Fig. 6).

## Discussion

We have uncovered here a molecular mechanism of memory mediated by H3K27me3 histone modification in *A. thaliana*, which is similar to that in *Drosophila melanogaster*^23, 24^. These data thus demonstrate that epigenetic memory mechanisms are conserved in species from two kingdoms of life. In particular, our results show that H3K27me3 demethylases specifically control the histone modifications of a few key target genes to maintain heat memory under a lab condition. We also determined that the H3K27me3-demethylase-mediated memory mechanism we observed in lab conditions functions under fluctuating field temperature conditions. Moreover, orthologs of key heat-memory genes, such as *Ahg474482* (*HSP17*.*6C*), *Ahg489869* (*HSP22*), and *Ahg945273* (*HSP21*), are highly expressed in field-grown *Arabidopsis halleri* during summer^25^. These results may suggest that those demethylases play a key role in heat acclimation and *HSP* regulation in the field. Interestingly, more genes were differentially expressed in *jmjq* mutants under fluctuating field temperature conditions than under a lab condition. Furthermore, differences in gene expression and histone modification in the wild type and the *jmjq* mutant under fluctuating field temperature conditions were more prevalent than those under a lab condition. Thus, additional memory pathways probably exist. Further support for our findings comes from a recent study of the acquisition of freezing tolerance in response to rapid and gradual decreases of temperature^26^. During evolution, plants may well have developed more than one system to respond and memorize rapid and gradual changes in temperature. Together with H3K27me3, other histone modifications or epigenetic changes, such as H3K36me3, DNA methylation and chromatin structure alterations, might also contribute to heat-stress memory^18, 27, 28^. Alternatively, H3K27me3 demethylases may remove repressive histone marks repeatedly in response to each and every recurring heat stress under fluctuating field temperature conditions. Moreover, those two possibilities are not mutually exclusive. Our findings, combined with such potential future discoveries, will further elucidate the mechanisms of plant adaptation and memory.

## METHODS

### Plant materials and growth conditions

*Arabidopsis thaliana* plants were grown at 22°C under continuous light unless specified. Culture medium consisted of half-strength Murashige and Skoog (MS) salts (Nacalai Tesque) and 0.8% agar (Nacalai Tesque) at pH 5.6 was used for all experiments except for crossing and transformation. For mature plant materials, seeds were sown in pots containing vermiculite and Metro-Mix. The following plant lines were previously described: *jmj30-2*^4^, *jmj32-1*^4^, and *jmj30-2 jmj32-1*^4^ mutants and *pJMJ30::JMJ30-HA*^4^; *elf6-3*^29^ and *ref6-1*^29^; and *hsp22*^15^. The *hsp17*.*6c-1* (SALK_56872) mutant line was obtained from the Arabidopsis Biological Resource Center. All plants were in the Columbia background. Genotyping primers are listed in Supplementary Table 13.

### Plasmid construction and plant transformation

For the *GUS* constructs, the genomic regions of the *HSP22, HSP17*.*6C*, and *HSP21* loci, including sequence upstream of the translational start site, and coding regions, and excluding the translation termination codon, were amplified by PCR using PrimeSTAR GXL DNA polymerase (Takara) and gene-specific primer sets. The resulting PCR products were subcloned into the pENTR/d-TOPO vector (Thermo Fisher Scientific). After sequences were confirmed with an ABI3130 sequencer (ABI), the insert fragments were transferred into the pBGWF7 destination vector^30^ using LR clonase (Thermo Fisher Scientific). All the *GUS* constructs were transformed into wild-type plants by floral dip using *Agrobacterium tumefaciens* (GV3101)^31^. T_1_ plants grown on soil were treated with the nonselective herbicide Basta. T_2_ seeds were harvested from Basta-resistant plants. *GUS* expression testing with and without heat treatment was performed using at least 15 independent T_2_ plants. Representative lines were used for further analysis. Cloning primers are listed in Supplementary Table 13.

For the overexpression construct, the *HSP17*.*6C* cDNA was amplified by PCR using PrimeSTAR GXL DNA polymerase (Takara) and gene-specific primer sets. cDNA prepared from *A. thaliana* Columbia was used as a template. The resulting PCR products were subcloned into pENTR/d-TOPO vector (Thermo Fisher Scientific). After sequences were confirmed with an ABI3130 sequencer (ABI), the insert fragments were transferred into the pB2GW7.0 destination vector^30^ using LR clonase (Thermo Fisher Scientific). The construct was transformed into *jmjq* plants by floral dip using *Agrobacterium* (GV3101)^31^. T_1_ plants grown on soil were treated with Basta. T_2_ seeds were harvested from Basta-resistant plants. Expression levels of *HSP17*.*6C* were examined using more than 15 independent T_2_ plants. Representative lines were used for further analysis. Cloning primers are listed in Supplementary Table 13.

For the *pER8::JMJ30* construct, *JMJ30* cDNA was amplified by PCR using gene-specific primer sets with restriction enzyme sequences. cDNA prepared from *Arabidopsis* Columbia was used as a template. The amplified DNA fragment was Gateway-cloned into pENTR/d-TOPO vector (Thermo Fisher Scientific). After sequencing using vector-specific primers, the fragment was introduced into the pER8 vector^21^. The construct was transformed into wild-type plants by floral dip using *Agrobacterium* (GV3101)^31^. T_1_ plants were grown on MS medium supplemented with hygromycin for transgenic selection. More than 20 independent inducible lines were further screened with phenotypic and expression analyses. Representative lines were used for further analysis. Cloning primers are listed in Supplementary Table 13.

### Heat stress treatment

*Arabidopsis* seeds were sterilized with 1 ml of 100% bleach in a 1.5-ml tube for <1 min. After being washed at least three times with sterilized water, seeds were sown on half-strength MS medium containing 0.8% agar. The plates were kept at 4°C for 5–7 days to synchronize seed germination. Plates were then moved into growth chamber at 22°C under continuous light. After 24 hrs, radicle tip emergence (germination) was observed regardless of genotype. When required, seedlings were subjected to heat-stress treatment. For heat-stress treatment in laboratory conditions, a PERSONAL-11 water bath shaker (TAITEC) was used^32^. Plates were placed into re-sealable zipper storage bags (S.C. Johnson & Son) and submerged in the water bath. Two kinds of stress were used: acclimation treatment (37°C for 20 min) and heat-shock treatment (43.5°C for 60 min) were performed on 4-day-old (96 hrs after germination) and 7-day-old (168 hrs after germination) seedlings, respectively. After each heat treatment, seedlings were returned to the growth chamber and allow to recover for further experiments.

To grow plants under fluctuating temperature conditions, seeds were prepared as described above. Plants were grown for 4 days after germination in a growth chamber at 22°C under continuous light. Then, the plants were moved and grown in a SGCmini growth chamber (Clockmics inc.) under fluctuating temperature conditions. The conditions used followed the actual environmental data for Tsubetsu (lat. 43°42.1’ N. and long. 144°2.0’ E.) from August 21 to 27, 2012; Sendai (lat. 38°15.7’ N. and long. 140°53.8’ E.) from July 13 to 19, 2018; Nara (lat. 34°41.6’ N. and long. 135°49.6’ E.) from August 4 to 10, 2018; Miyazaki (lat. 31°56.3’ N. and long. 131°24.8’ E.) from August 3 to 9, 2017; and Naha (lat. 26°12.4’ N. and long. 127°41.2’ E.) from July 18 to 24, 2015. Past temperature data from five cities at different latitudes in Japan were obtained from the Japan Meteorological Agency (https://www.data.jma.go.jp/obd/stats/etrn/index.php). The temperature in the SGCmini growth chamber was changed for 6 days following the pattern for a given city. Temperatures recorded in the SGCmini growth chamber every hour are listed in Supplementary Table 9.

### Estradiol treatment

For β-estradiol treatment, the compound was dissolved in dimethyl sulfoxide (DMSO) just prior to use. For mock treatment, the same amount of DMSO was used as control. Plants were grown on MS medium without β-estradiol and then transplanted onto MS medium with 10 μM β-estradiol^21^ using ethanol-sterilized forceps in a hood. Once we started treatment, plants were transplanted every 2 days.

### Phenotypic analyses

For all phenotypic analyses, plants to be directly compared were grown side by side at the same density per plate to minimize potential microenvironmental differences in the growth chamber. To determine the survival rate, phenotypic strength in the wild type and each mutant were categorized into three different classes^33^. If plants were vigorous and looked entirely green, those plants were categorized as “normal”. If plants were largely pale, those were counted as “partially damaged”. Dead plants were categorized as “perished”. *n* > 138. At least three independent experiments were performed, and similar results were obtained. Representative plate images were photographed with a Nikon D750 camera. A chi-squared (χ^2^) test, followed by post-hoc test, was performed through IBM SPSS Statistics 26 (IBM).

For chlorophyll measurement, total chlorophyll levels were measured using *N,N*′-dimethylformamide (DMF) extraction and spectrophotometric quantification^34^. Five seedlings were placed into a 2-ml tube containing 1 ml of DMF. The experiments were repeated more than 5 times. *n* > 5. Tubes were incubated overnight at 4°C. The absorbance at 646.8 and 663.8 nm was measured in 1.00-cm cuvettes on spectrophotometer (IMPLEN NanoPhotometer P-Class). At least three independent experiments were performed, and similar results were obtained. Total chlorophyll was calculated as reported previously^34^ (Chl *a* + *b* (μM) = 19.43 *A*646.8 + 8.05 *A*663.8). Normal distribution was verified by the Kolmogorov-Smirnov test for all analyses. Statistical significance was computed using a one-way ANOVA test followed by post-hoc Tukey’s HSD test (https://astatsa.com/OneWay_Anova_with_TukeyHSD/).

For fresh weight assay, 10-day-old seedlings were used. Ten *Arabidopsis* seedlings were harvested in a 2-ml tube and measured with an analytical balance^35^ (Mettler Toledo XS104). *n* > 11. At least three independent experiments were performed, and similar results were obtained. Normal distribution was verified by the Kolmogorov-Smirnov test for all analyses. Statistical significance was computed using one-way ANOVA test followed by post-hoc Tukey’s HSD test (https://astatsa.com/OneWay_Anova_with_TukeyHSD/).

For leaf measurement, 5th true leaves from 10-day-old seedlings were used. Leaves were dissected using forceps and surgical scissors. Images were taken with a Nikon D750 camera and were used to measure leaf area with ImageJ software (http://rsb.info.nih.go.ij/). To quantify cell size and number, leaves were nicked at the edges and the resulting samples were placed into fixation solution and placed under vacuum for 20 min for infiltration. The resulting leaf tissue samples were transferred into clearing solution and kept for at least 16 hrs at room temperature. The samples were then mounted onto microscope slides with one or two drops of clearing solution, and imaged using an AxioScope A1 microscope (Zeiss). The cell area was measured with ImageJ software (http://rsb.info.nih.go.ij/). Leaf and cell areas were subsequently used to calculate cell numbers^36^. Statistical significance was computed using one-way ANOVA test followed by post-hoc Tukey’s HSD test (https://astatsa.com/OneWay_Anova_with_TukeyHSD/).

### Scanning Electron Microscopy (SEM)

For SEM, the aboveground parts of 10-day-old seedlings were placed in FAA (45% EtOH, 2.5% formaldehyde, and 2.5% acetic acid), placed under vacuum until the tissues sank into the fixative, and left at least 16 hrs at room temperature. Tissues were then transferred successively through a gradient series of ethanol solutions (50%, 60%, 70%, 80%, 90%, 95%, 100% × 2) in water for 20 min each, and then a gradient series of acetone (25%, 50%, 75%, 95%, 100% × 2) in ethanol for 30 min each. Then, the tissues were critical-point-dried with liquid CO_2_ in an EM CPD300 critical-point dryer (Leica Microsystems) and gold-coated with E-1010 (Hitachi) prior to SEM imaging. The tissues were imaged under an S-4700 SEM (Hitachi) with an accelerating voltage of 15 kV. At least five shoot apical meristems for each genotype and treatment were observed, and representative images are shown.

### RNA analyses

For RNA extraction, total RNA was extracted from whole seedlings using the RNeasy Plant Mini Kit (Qiagen). About 100 mg of plant tissues was harvested, immediately frozen in liquid nitrogen, and kept at −80 °C until use. The tissues were ground to a fine powder with ice-cold mortar and pestle. RNA extraction was performed following the manufacturer’s instructions. An RNase-Free DNase Set (Qiagen) was used to remove genomic DNA. Concentration was measured by spectrophotometer (IMPLEN NanoPhotometer P-Class).

For qRT-PCR, cDNA was synthesized from 100 ng of RNA using a PrimeScript 1^st^ strand cDNA Synthesis Kit (Takara). The resulting cDNA was quantified with a LightCycler 480 (Roche) using FastStart Essential DNA Green Master mix (Roche). The signals were normalized against the internal control gene *UBIQUITIN CONJUGATING ENZYME 21*^37^ (*UBC21*; *AT5G25760*). Three independent experiments were performed. Each result is shown by jitter plot. Statistical significance was computed using Student’s *t*-test. Primers for qRT-PCR are listed in Supplementary Table 13.

For RNA-seq, libraries were prepared as reported previously^38^ and sequenced by HighSeq 2500 using 50-base-pair single-end mode (Illumina). Mapping was conducted using the *Arabidopsis thaliana* reference genome (TAIR10). The read count for each gene was calculated by RSEM^39^. After normalization, FDR and FC were calculated using the edgeR package for R^40^. Genes with false discovery rate (FDR) < 0.05 in each comparison were identified as differentially expressed genes. The agriGO web-based tool and database (http://bioinfo.cau.edu.cn/agriGO/) was used for a Gene Ontology (GO) term enrichment analysis^41,42^. Heatmaps and *k*-mean clustering graphs were generated with MeV (http://mev.tm4.org/#/welcome). The sequence data were deposited into the DNA Data Bank of Japan (DRA008818, DRA009425).

### Trypan blue staining

To identify dead cells, trypan blue staining was performed as previously described with minor modification^43^. Cotyledons were collected into 1.5-ml tubes containing 0.05% trypan blue (Nacalai Tesque). The tubes were then boiled for 1 min to stain dead cells. After staining, tissues were transferred into chloral hydrate solution and kept at least 6 hrs. The resulting tissues were placed on glass slides, mounted in a drop of 50% glycerol, and immediately observed under an AxioScope A1 microscope (Zeiss) equipped with an AxioCam ERc 5s camera (Zeiss) and analyzed using the ZEN2 software (Zeiss). At least five cotyledons were observed, and representative images are shown.

### DAB staining

To observe reactive oxygen species (ROS) accumulation, H_2_O_2_ staining was conducted as previously described with minor modification^44^. Cotyledons were collected into 1.5-ml tubes and stained using the peroxidase stain DAB Kit (Nacalai Tesque) for 2 hrs in darkness with gentle shaking. After staining, tissues were transferred to fresh tubes containing bleach solution (60% ethanol, 20% acetic acid, 20% glycerol) and boiled for 15 min to decolorize chlorophyll. The resulting tissues were placed on glass slides, mounted on a drop of 50% glycerol, and immediately observed under an AxioScope A1 microscope (Zeiss) equipped with an AxioCam ERc 5s camera (Zeiss) and analyzed using the ZEN2 software (Zeiss). More than five cotyledons were observed, and representative images are shown.

### GUS expression analysis

For GUS staining, tissues were kept in 90% acetone for 20 min to infiltrate solution. Tissues were rinsed with GUS staining buffer without 5-bromo-4-chloro-3-indolyl-β-D-glucuronic acid (X-Gluc) three times and put into GUS staining buffer with X-Gluc. They were placed under vacuum until the tissues sank, and then left overnight or for two nights, depending on the expression levels. After GUS staining, chlorophyll was removed by allowing tissues to sit in 70% EtOH for at least 1 week. Representative images were taken under an AXIO Zoom.V16 (Zeiss) microscope.

To quantify GUS activity, a MUG assay was performed as previously described with minor modification^45^. Ten-day-old seedlings were harvested, immediately frozen in liquid nitrogen, and kept at −80 °C until use. Tissues were homogenized in extraction buffer and protein solution was obtained after removal of debris by centrifugation. Reaction buffer containing 4-methylumbelliferyl-β-D-glucuronide (4-MUG) was mixed with the resulting protein solution. Reactions were stopped after 0 and 60 min incubation at 37°C by adding stop buffer. 4-MU fluorescence was measured with a Tristar^2^-LB942 microplate reader (BERTHOLD) using an excitation and emission wavelengths of 365 and 455 nm, respectively. Protein amounts were determined using a Qubit4 Fluorometer and Qubit Protein Assay kit (Thermo Fisher Scientific). GUS activity was calculated in (nmol 4-MU) min^-1^ (mg protein)^-1^. Statistical significance was computed using one-way ANOVA test followed by post-hoc Tukey’s HSD test (https://astatsa.com/OneWay_Anova_with_TukeyHSD/).

### Chromatin immunoprecipitation (ChIP)

For ChIP-qPCR, ChIP was carried out as described previously^46^. For each sample, 100–300 mg of seedlings were fixed with 1% formaldehyde for 15 min. After quenching of formaldehyde with glycine for 5 min, tissues were frozen in liquid nitrogen and kept at −80°C until use. Tissues were ground to a fine powder with an ice-cold mortar and pestle. Using nuclei extraction buffer, chromatin was isolated from a nuclear extract. Fragmentation was conducted using an Ultrasonic Disruptors UD-201 sonicator (TOMY). After preclearing, antibodies were added and the mixtures were rotated overnight at 4°C. H3K4me3 (ab8580; Abcam), H3K27me3 (ab6002; Abcam), H3 antibodies (ab1791; Abcam), and HA (12CA5; Roche) were used. For immunoprecipitation to capture DNA-protein complexes, Dynabeads Protein A or G (Thermo Fisher Scientific) were used. Beads were washed and DNA was eluted from beads overnight at 65°C. The resulting DNA was purified using QIAquick PCR Purification Kit (Qiagen). DNA was quantified with a LightCycler 480 (Roche) using FastStart Essential DNA Green Master mix (Roche). The ratio of ChIP over input DNA (% Input) was compared based on the reaction threshold cycle for each ChIP sample compared to a dilution series of the corresponding input sample. Three independent experiments were performed. Each result was shown by jitter plot. Statistical significance was determined by one-way ANOVA followed by post-hoc Tukey’s HSD test for multiple-pair comparisons or a two-tailed Student’s *t*-test for single-pair comparisons. Primers for qPCR are listed in Supplementary Table 13.

ChIP-seq was performed as previously described with minor modifications^47^. 1.5 g of seedling tissues were frozen in liquid nitrogen and kept at −80°C before use. Tissues were ground to a fine powder with an ice-cold mortar and pestle and post-fixed in nuclei isolation buffer for 10 min. Glycine was added and kept for 5 min at room temperature. After removal of debris with a Miracloth (Merck), chromatin was dissolved into ChIP dilution buffer and sonication was conducted by S2 sonicator (Covaris). Chromatin and antibody were mixed and rotated overnight at 4°C. Antibodies were described above. Dynabeads M280 Sheep anti-mouse IgG or Dynabeads Protein G (Thermo Fisher Scientific) were used for immunoprecipitation. Beads were washed, and chromatin was eluted by ChlP direct elution buffer. The resulting chromatin was incubated overnight at 65°C to reverse crosslinking. After digesting RNase and Proteinase K, ChIP DNA was purified with the Monarch PCR & DNA Cleanup Kit (NEB). Libraries were prepared using ThruPLEX DNA-seq Kit (Rubicon Genomics) according to the manufacturer’s instructions. Dual size selection was performed using Agencourt AMpure XP beads (Beckman Coulter). The libraries were pooled and sequenced by Next-Seq 500 (Illumina). Two independent biological replicates were analyzed for each genotype. The sequence data were deposited into the DNA Data Bank of Japan.

Prior to mapping of reads onto the *Arabidopsis thaliana* TAIR10 genome, trimming and filtering of reads were conducted. Bowtie^48^ with -m 1 -best parameters was used to control multi-reads and SAM files were obtained. The SAM files were converted to sorted BAM files using SAMtools^49^. The resulting BAM files were then converted to BED files through BEDTools^50^. To extend the 5’ end of reads toward the 3’ direction, the slop function in the BEDTools was utilized. Mapping were performed on the NIG supercomputer at the ROIS National Institute of Genetics. The resulting reads were counted using the coverage function in the BEDTools. Hypermethylated regions of H3K27me3 and hypomethylated regions of H3K4me3 in *jmjq* compared to WT were identified using a fold enrichment threshold of 1.5 by R. To visualize binding peaks, Integrative Genome Viewer^51^ was used. ngs.plot was used to generate metablot and heatmap for ChIP-seq data^52^.

## Note

Any Supplementary Information and Source Data files are available in the online version of the paper.

## ACKNOWLEDGMENTS

We thank Akie Takahashi, Hiroko Egashira, Kyoko Sunuma, Mayumi Nara, Mikiko Higashiura, Taeko Kawakami, and Yuka Kadoya for technical assistance; Kei Hiruma and Masanori Izumi for suggestions about the cell death and ROS accumulation experiments; and Sachi Ando for helping statistical analysis by SPSS and checking the draft of this manuscript. Computations were partially performed on the NIG supercomputer at the ROIS National Institute of Genetics. This work was supported by a grant from the Japan Science and Technology Agency ‘PREST’ (JPMJPR15QA), a JSPS KAKENHI Grant-in-Aid for Scientific Research on Innovative Areas (No. 18H04782), a JSPS KAKENHI Grant-in-Aid for Scientific Research B (No. 18H02465), a Grant-in-Aid for challenging Exploratory Research (No. 19K22431), and a grant from the Mishima Kaiun Memorial Foundation to N.Y., a grant from the Japan Science and Technology Agency ‘PREST’ (JPMJPR17Q1) to S.I., a JSPS KAKENHI Grant-in-Aid for Scientific Research C (No. 15H05955) to T.S., a JSPS KAKENHI Grant-in-Aid for Scientific Research on Innovative Areas (No. 15H05963) to T.K., a Japan Science and Technology Agency ‘CREST’ (JPMJCR15O2) grant to A.J.N., a JSPS KAKENHI Grant-in-Aid for Scientific Research on Innovative Areas (No. 17H06478) to A.S., and a JSPS KAKENHI Grant-in-Aid for Scientific Research on Innovative Areas (No. 19H04865, 20H04888), a JSPS KAKENHI Grant-in-Aid for Scientific Research A (No. 15H02405, 20H00470), and a Grant-in-Aid for challenging Exploratory Research (No. 18K19342) to T.I. The RNA-seq and ChIP-seq data have been deposited in the DDBJ database (DRA008818, DRA009425).

## Author contributions

Conceptualization: N.Y. and T.I. (lead) and all other authors (supporting); data curation: N.Y.; formal analysis: N.Y., S.M., and K.Y.,; funding acquisition: N.Y., S.I., T.S., T.K., A.J.N., A.S., and T.I.; investigation: N.Y. (lead) and S.M., K.Y., M.S., K.H., M.K., Y.K., S.I., T.S., E.-S.G., and T.T. (supporting); project administration: N.Y.; software: N.Y. (lead) and M.S., K.H., M.K., S.I., T.S., T.T., A.J.N., and A.S.; supervision: N.Y. and T.I.; validation: N.Y.; visualization: N.Y.; writing: N.Y. (original draft) and all authors (review and editing).

