## Supplemental file for "Removal of repressive histone marks creates epigenetic memory of recurring heat in *Arabidopsis*"

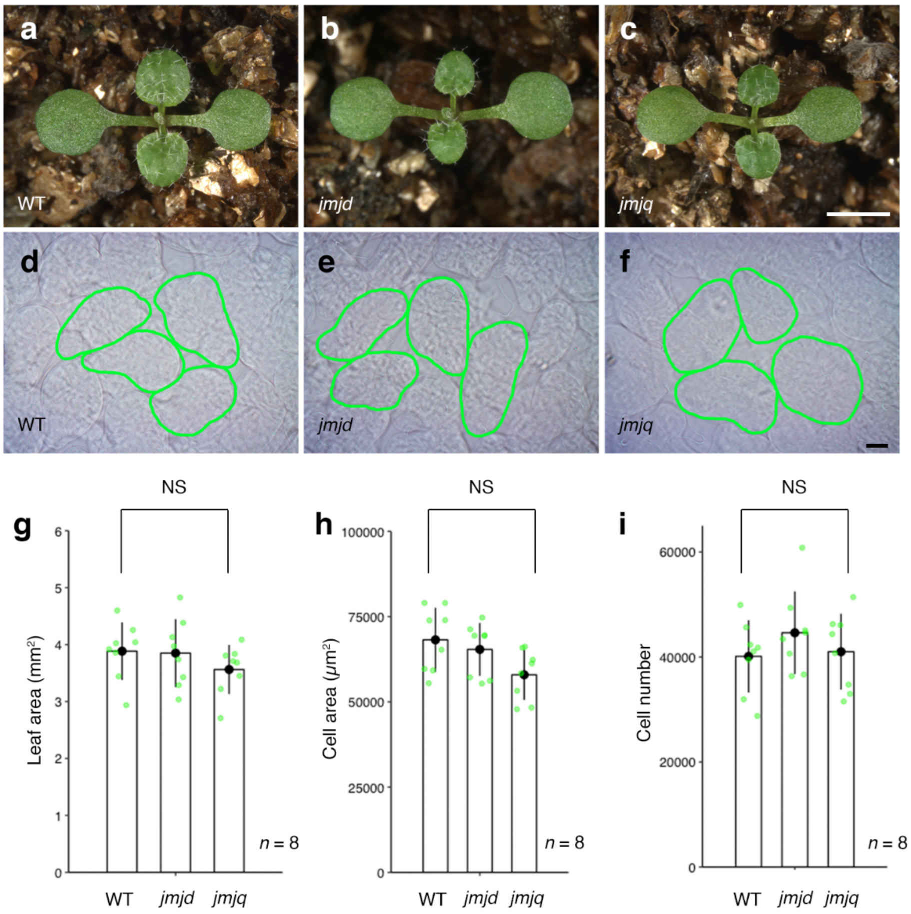


**Extended Data Fig. 1 Leaf phenotypes in 10-day-old wild-type, *jmjd*, and *jmjq* seedlings. a**–**c**, Seedlings of the wild type (a) and *jmjd* (b) and *jmjq* mutants (c) at 10 days after germination. Bar = 1 mm. **d**–**f**, Palisade cells in the cotyledons of wild type (d), *jmjd* (e), and *jmjq* (f). Representative cells are marked in green. Bar = 20 μm. **g**–**i**, Leaf area (g), cell area (h), and projected subepidermal palisade cell number (i) of wild type, *jmjd*, and *jmjq* at 10 days after germination. NS, nonsignificant based on one-way ANOVA test. $p < 0.05$. No significant differences in leaf area, cell area, or cell number were observed in these three mutants grown under control condition. These result suggest that acclimation defects seen in the *jmjq* mutant are not caused by morphological changes.

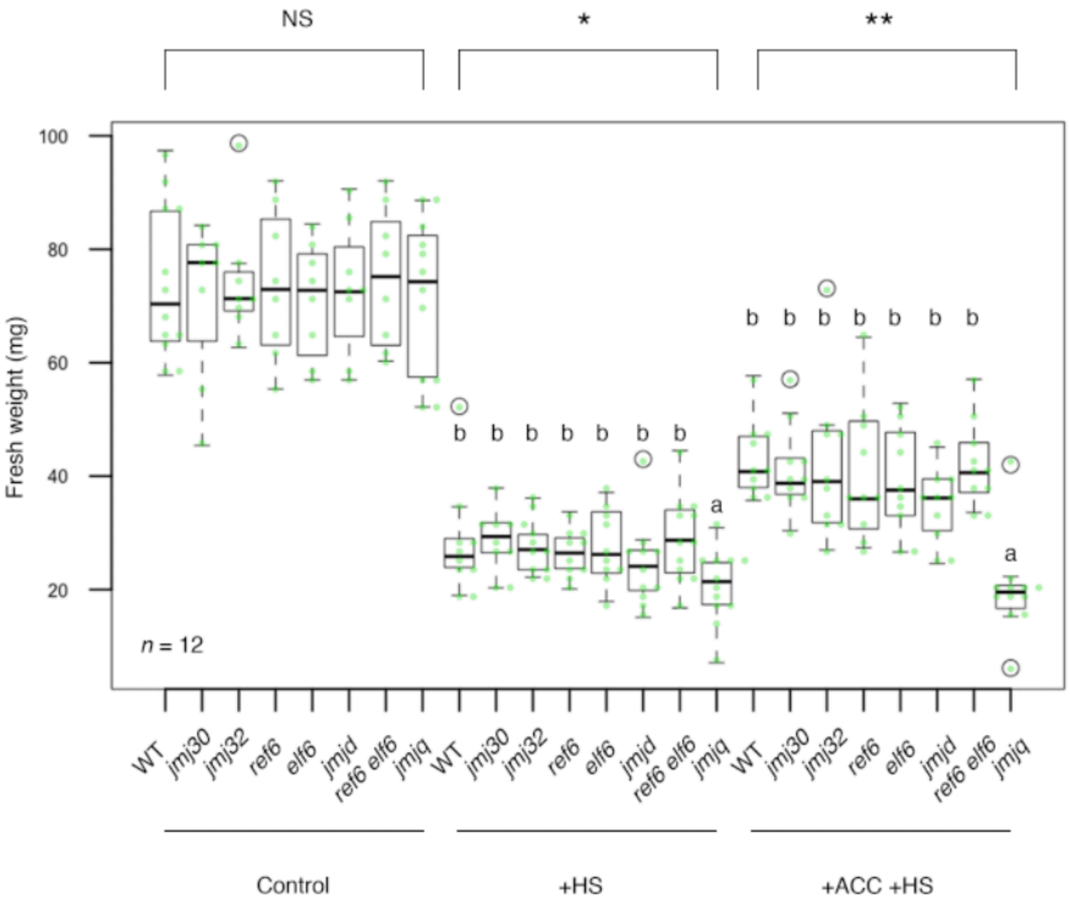


**Extended Data Fig. 2 Heat acclimation phenotype in *jmjq* mutants.** Quantification of seedling fresh weights. Light green jitter dots and white circles represent the fresh weight from each sample and from statistical outliers, respectively. Asterisks indicate significant differences based on one-way ANOVA test. *$p < 1.0 \times 10^{-3}$, $p < 1.0 \times 10^{-4}$. Different letters indicate significant differences, while the same letters indicate non-significant differences based on post-hoc Tukey's HSD test. $p < 0.05$. NS, nonsignificant.

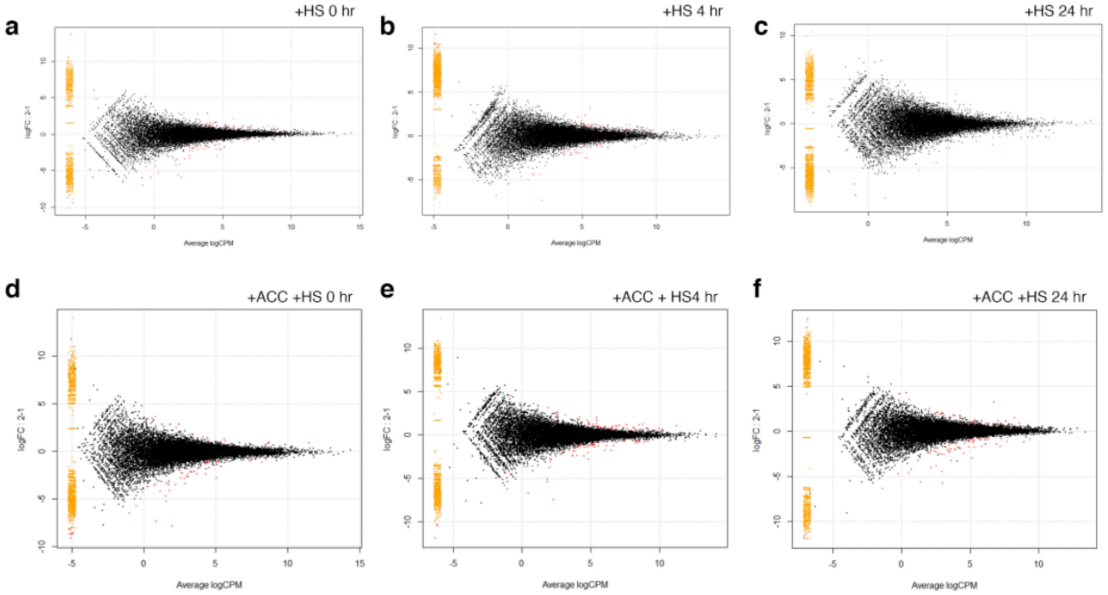


**Extended Data Fig. 3 MA plots of the log fold change of all genes. a–f**, RNA-seq results under basal thermotolerance condition (a-c) and heat-stress memory condition (d-f). Timing for RNA-seq, is shown in Fig 2a. The MA plots represent each gene with a dot. The *x* axis is the average log CPM over all genes; the *y* axis is the $\log_2$ fold change of normalized count between wild type and *jmjq* mutants. Genes with FDR <0.05 are shown in red. +HS 0 hr (a), +HS 4 hr (b), +HS 24 hr (c), +ACC +HS 0 hr (d), +ACC +HS 4 hr (e), and +ACC +HS 24 hr (f).

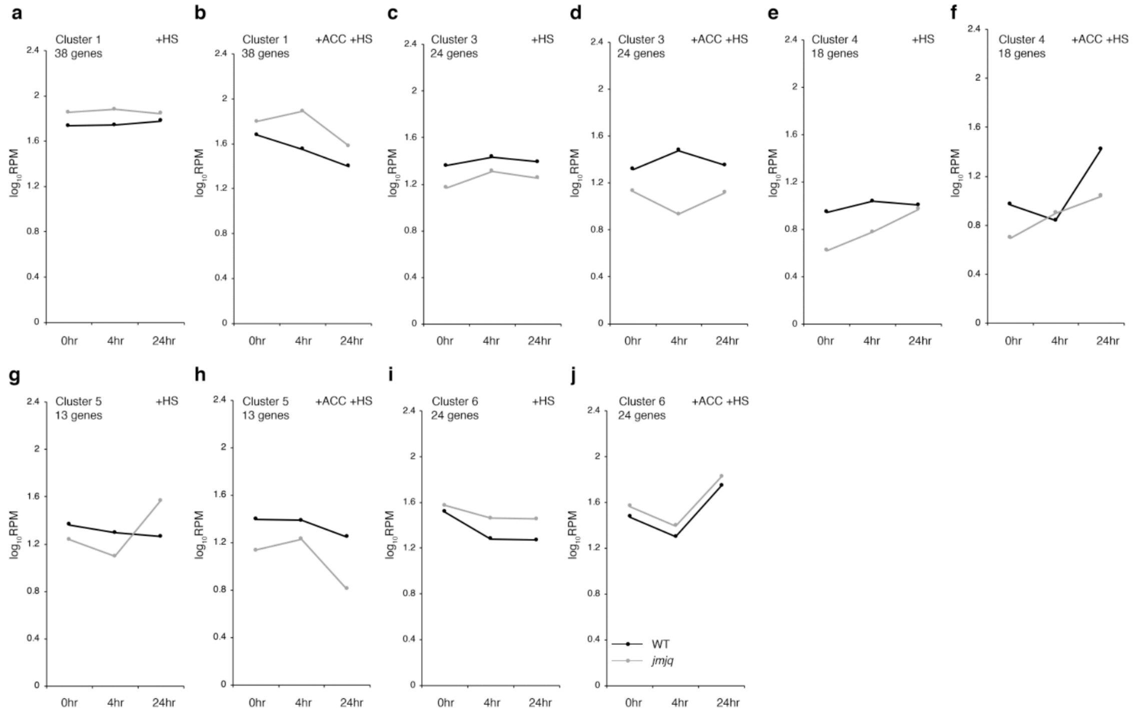


**Extended Data Fig. 4 Gene expression of from the six hierarchical clusters. a–j**, *k*-means clustering of genes differentially expressed between wild type and *jmjq* mutants. An optimal number of six clusters were identified using 142 differentially expressed genes in *jmjq* mutants grown under +ACC +HS conditions. Shown are cluster 1 +HS (a), cluster 1 +ACC +HS (b), cluster 3 +HS (c), cluster 3 +ACC +HS (d), cluster 4 +HS (e), cluster 4 +ACC +HS (f), cluster 5 +HS (g), cluster 5 +ACC +HS (h), cluster 6 +HS (i), and cluster 1 +ACC +HS (j). Two of these clusters comprise genes upregulated in the *jmjq* mutant: clusters 1 and 6 (62 genes). Four clusters comprise genes downregulated in the *jmjq* mutant: clusters 2, 3, 4, and 5 (80 genes). The graphs of cluster 2 are shown in Fig. 2. Since JMJs activate gene expression though demethylation of H3K27me3, 80 genes that were downregulated in the *jmjq* mutant were analyzed further.

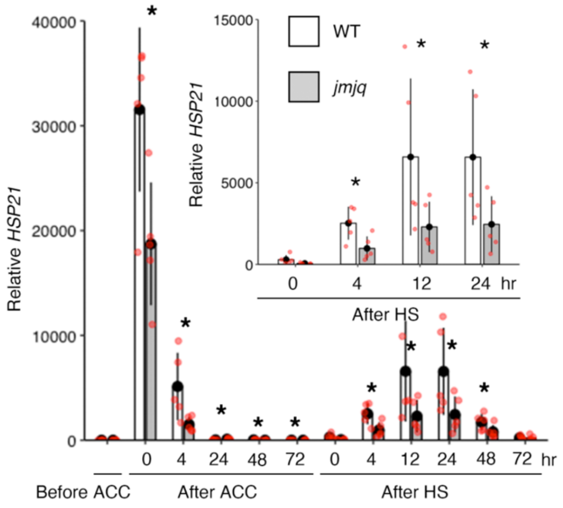


**Extended Data Fig. 5 *HSP21* expression with acclimation and heat shock.** qRT-PCR verification of the *HSP21* gene in the wild type and the *jmjq* mutant grown under the conditions shown in Fig. 2a. Red jitter dots represent expression level from each sample. Asterisks indicate significant difference at 0.05 levels between wild type and *jmjq* mutants at the same time pont based on Student's *t*-test. *HSP21* expression after HS was reduced in acclimatized *jmjq* mutants compared to acclimatized wild type.

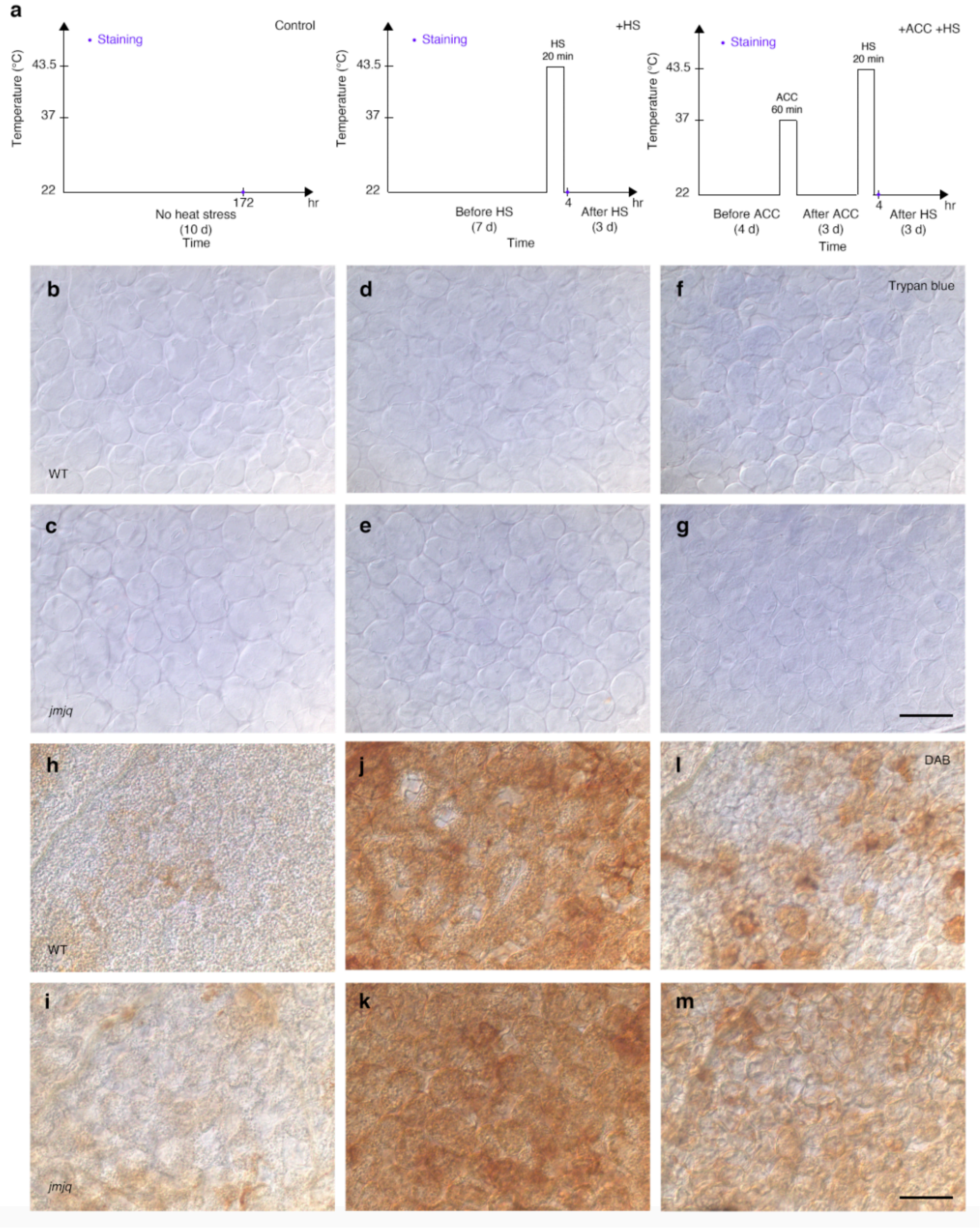


**Extended Data Fig. 6 Cell death and ROS accumulation after heat shock.**
**a**, Schematic representation of temperature conditions. Staining (navy dots) was conducted at 7 days after germination. Left, normal plant growth condition.

Center, basal thermotolerance condition. Staining was performed 4 hrs after heat shock. **b**–**m**, Trypan blue staining for the detection of cell death. (b and c) Wild type (b) and *jmjq* (c) grown under control condition. (d and e) Wild type (d) and *jmjq* (e) grown under +HS condition. (f and g) Wild type (f) and *jmjq* (g) grown under +ACC +HS condition. (h–m) DAB staining for the detection of $H_2O_2$ production. (h, i) Wild type (h) and *jmjq* (i) grown under control condition. (j and k) Wild type (j) and *jmjq* (k) grown under +HS condition. (l and m) Wild type (l) and *jmjq* (m) grown under +ACC +HS condition. Scale bars = 100 μm. The majority of cells are damaged after heat shock treatment. However, cells remained alive under all conditions at the time point shown in Extended Data Fig. 6a. These result suggest that the lack of expression of *HSP* genes right after the heat shock could be due to damage of cells.

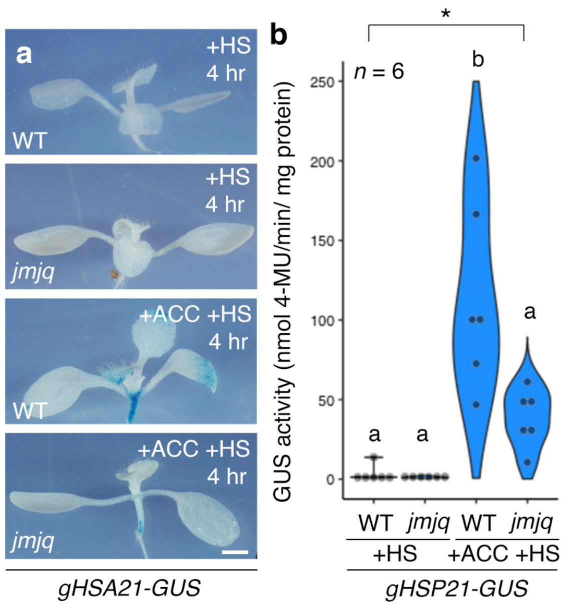


**Extended Data Fig. 7 HSP21-GUS accumulation in wild type and *jmjq* mutants. a**, gHSP21-GUS expression in wild type and *jmjq* mutants grown under the conditions shown in Fig. 2a. gHSP21-GUS was highly accumulated in acclimatized wild type after HS. *HSP21* expression was not detected in acclimatized *jmjq* mutants 4 hrs after HS. Scale bar = 1 mm. **b**, Quantification of GUS activity by MUG assay in the plants shown in Extended Data Fig. 7a. Asterisk indicates significant differences based on one-way ANOVA test ($p <$ 0.05). Different letters indicate significant differences, while the same letters indicate non-significant differences based on post-hoc Tukey's HSD test ($p <$ 0.05). NS, nonsignificant. $n$ = 5.

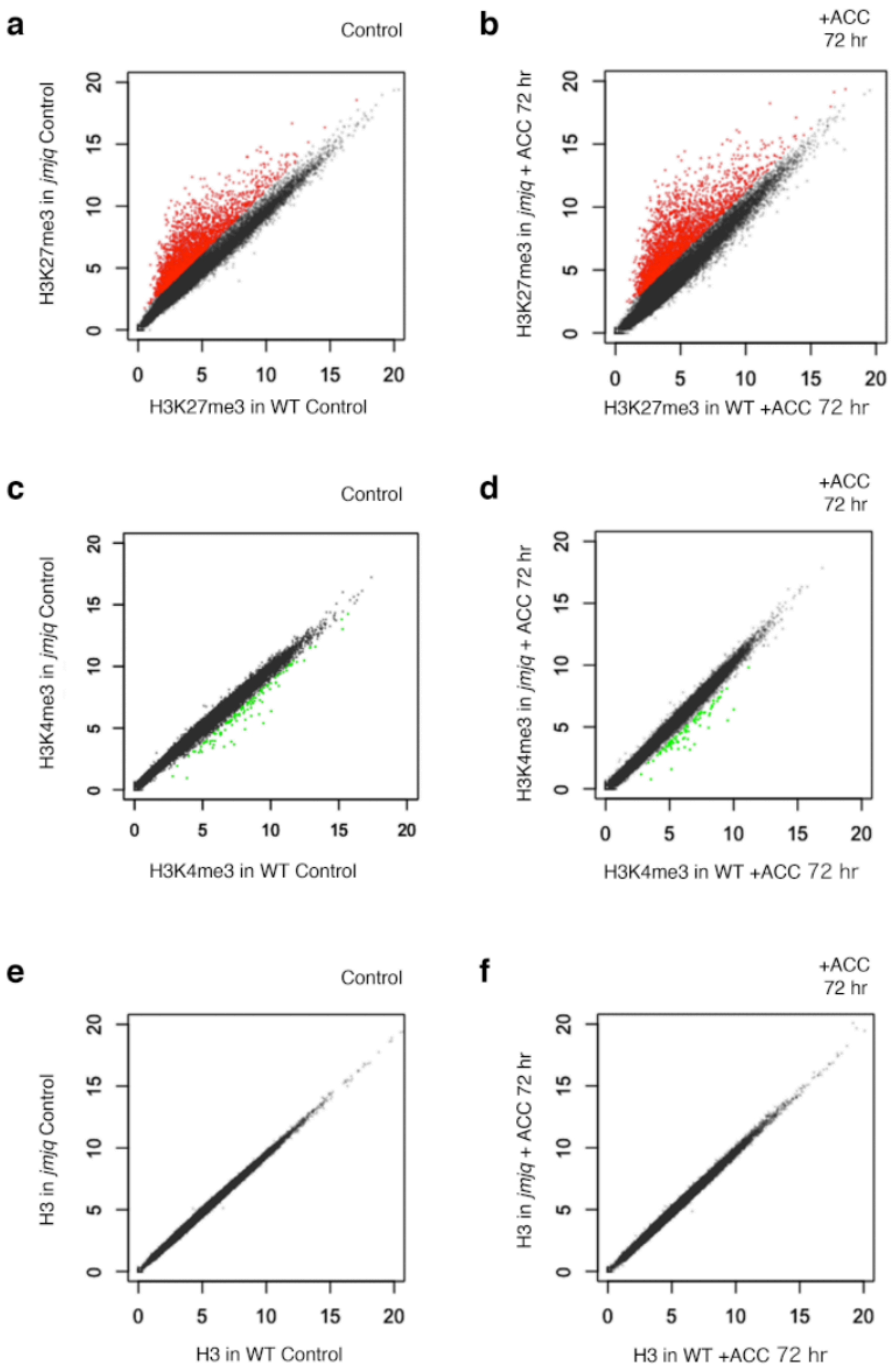


**Extended Data Fig. 8 Scatter plot of H3K27me3, H3K4me3, and H3 ChIP-seq a** and **b**, H3K27me3 levels determined by ChIP-seq in wild type and *jmjq* mutants without acclimation (a) and 3 days (72 hrs) after acclimation (b). **c** and **d**, H3K4me3 levels determined by ChIP-seq in wild type and *jmjq* mutants without acclimation (c) and 3 days (72 hrs) after acclimation (d). **e** and **f**, H3 levels determined by ChIP-seq in wild type and *jmjq* mutants without acclimation (e) and 3 days (72 hrs) after acclimation (f). Each dot represents the square root of the read counts per million mapped reads (RPM). Genes that were hyper H3K27 trimethylated in the *jmjq* mutant and hyper H3K4 trimethylated in wild type are shown in red and light green, respectively. Consistent with the biochemical role of JMJ in the removal of H3K27me3, more than 2000 genes were hypermethylated in the *jmjq* mutant. Hundreds of transcription start sites of genes were hypertrimethylated by H3K4me3 in wild type under two different conditions. No differences in H3 levels were observed between wild type and *jmjq* mutants with and without acclimation.

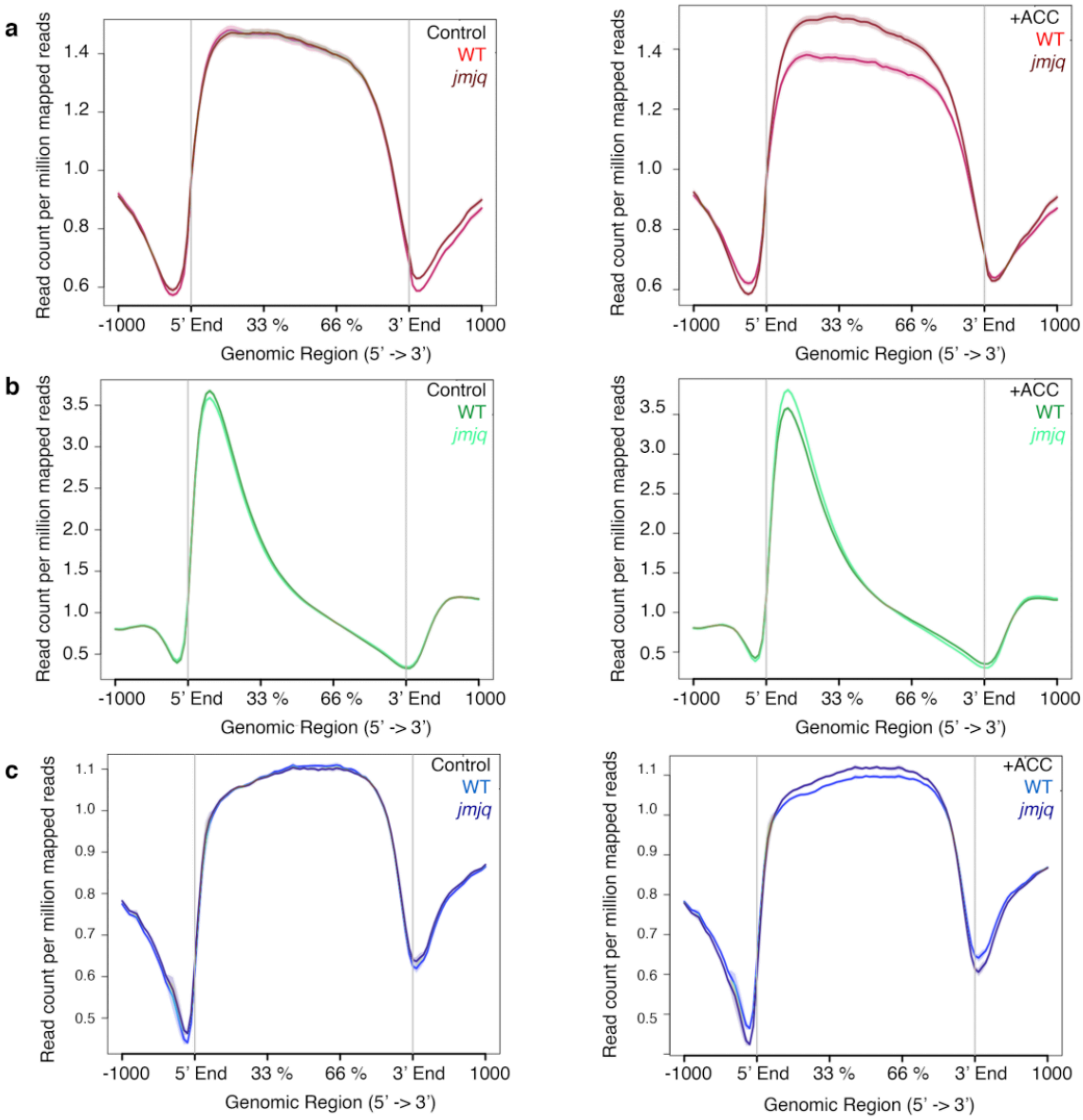


**Extended Data Fig. 9 Averaged profiles of H3K27me3, H3K4me3, and H3 around genes. a–c**, Metablots of H3K27me3 (a), H3K4me3 (b), and H3 (c) around genes in wild type and *jmjq* mutants without acclimation (left) and 3 days (72 hrs) after acclimation (right). Regardless of conditions or genotypes, H3K27me3 and H3K4me3 were observed in gene body and near transcription start sites. H3 enrichment in wild type and *jmjq* mutants with and without acclimation were also observed in gene bodies.

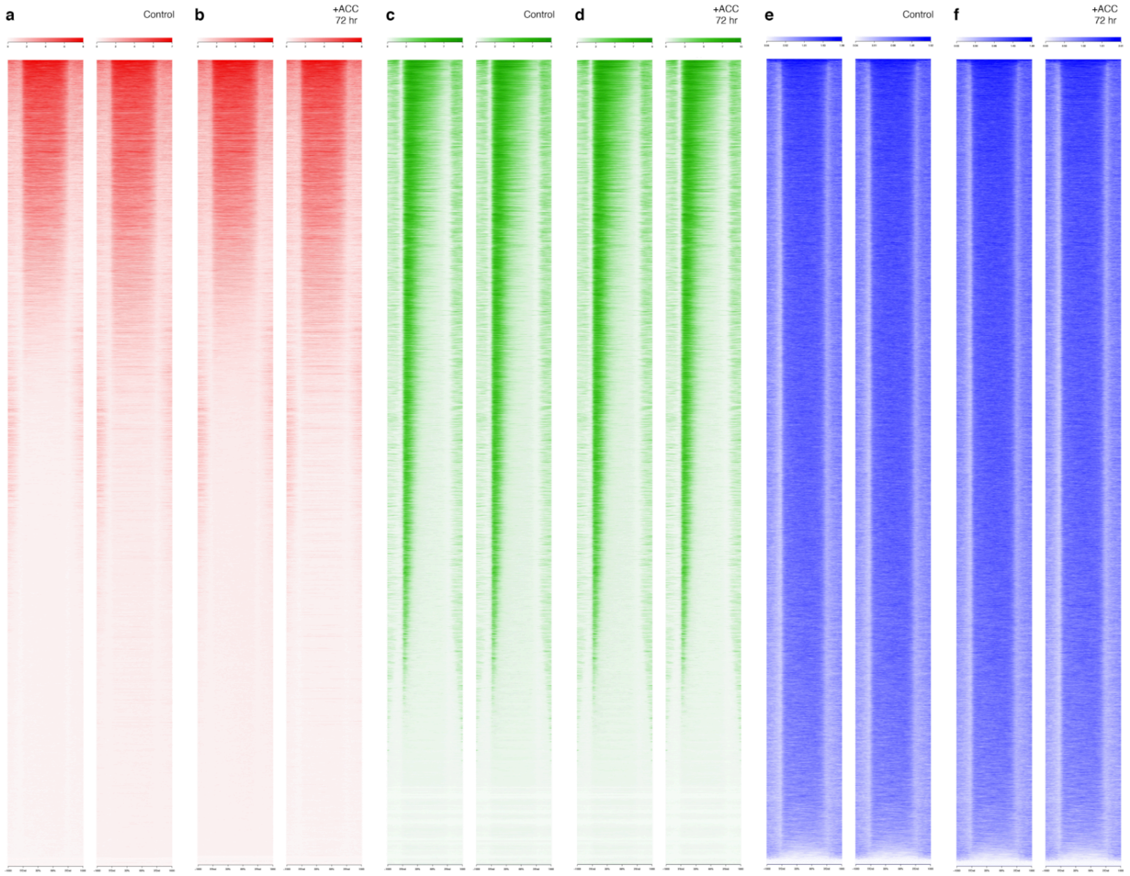


**Extended Data Fig. 10 Heatmaps of H3K27me3, H3K4me3, and H3. a** and **b**, Heatmaps of H3K27me3 around genes in wild type and *jmjq* mutants without acclimation (a) and 3 days (72 hrs) after acclimation (b). **c** and **d**, Heatmaps of H3K4me3 around genes in wild type and *jmjq* mutants without acclimation (c) and 3 days (72 hrs) after acclimation (d). **e** and **f**, Heatmaps of H3 around genes in wild type and *jmjq* mutants without acclimation (e) and 3 days (72 hrs) after acclimation (f).

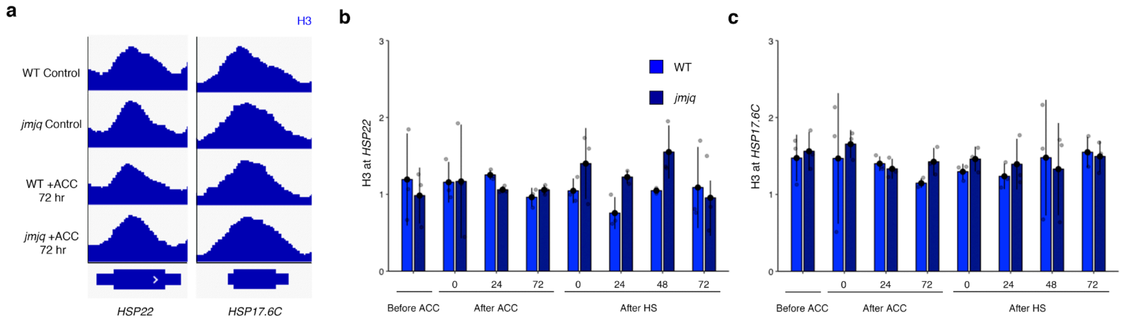


**Extended Data Fig. 11 Histone H3 levels at the *HSP22* and *HSP17.6C* loci in wild type and *jmjq* mutants. a**, H3 peaks detected by ChIP-seq at the *HSP22*, and *HSP17.6C* loci in wild type and *jmjq* mutants without acclimation (above) and 3 days (72 hrs) after acclimation (below). **b** and **c**, Histone H3 levels at *HSP22* (b) and *HSP17.6C* (c) determined by ChIP-qPCR in wild type and *jmjq* mutants. Grey jitter dots represent expression level from each sample. No difference in histone H3 signals was observed between wild type and *jmjq* mutants by Student's *t*-test. Taken together with the H3K27me3 and H3K4me3 results shown in Fig. 3, these data imply that it is histone modifications, rather than the locations of histones (or nucleosomes) along the DNA that are changed due to acclimation.

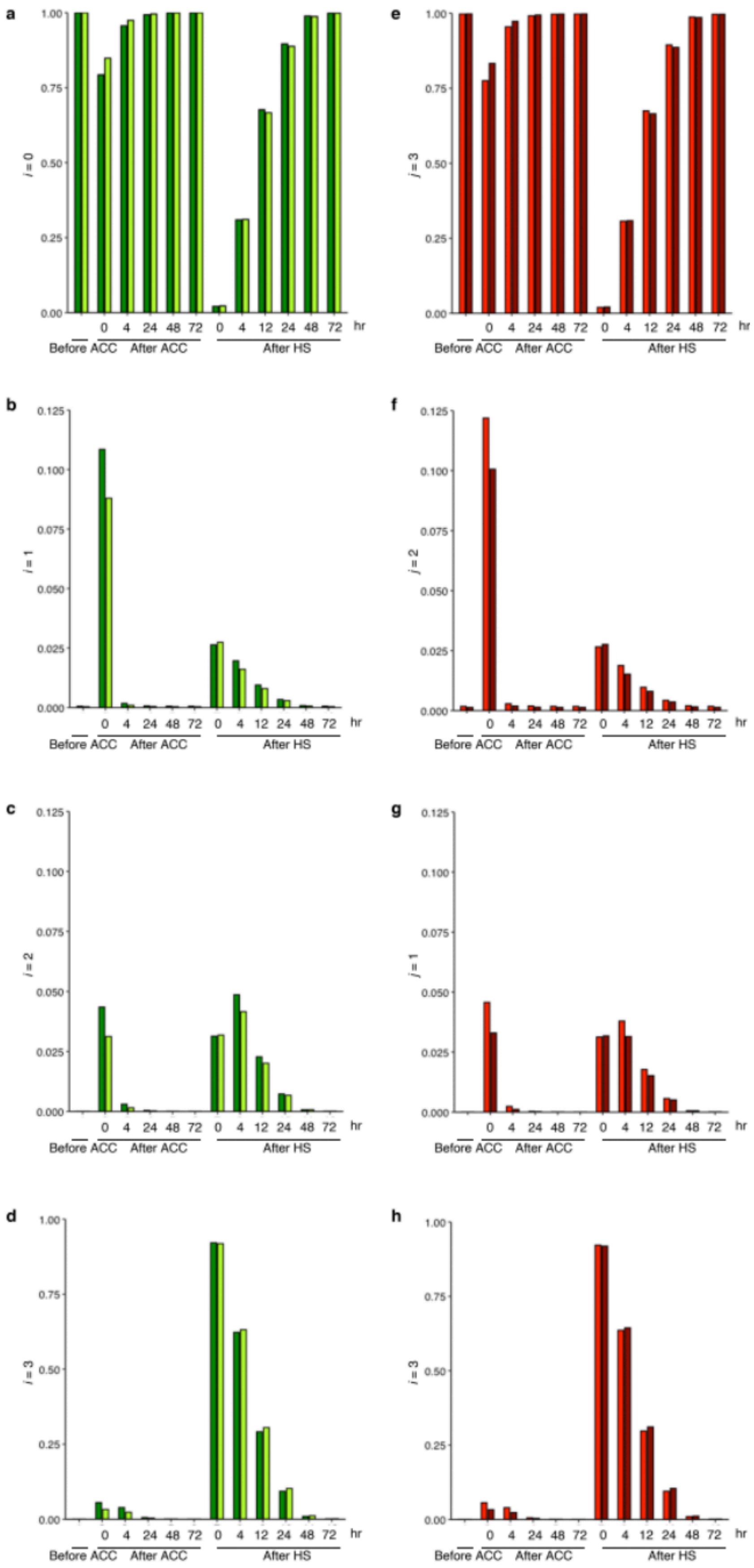
a
b
c
d
e
f
g
h
j = 0
j = 1
j = 2
j = 3
1.00
0.75
0.50
0.25
0.00
0.125
0.100
0.075
0.050
0.025
0.000
0 4 24 48 72 0 4 12 24 48 72 hr
Before ACC
After ACC
After HS

**Extended Data Fig. 12 The memory effects of having multiple actively modified nucleosomes. a**–**d**, Change in fraction of cells that have 0 (a), 1 (b), 2 (c), and 3 (d) actively modified nucleosomes. **e**–**h**, Change in fraction of cells that have 3 (e), 2 (f), 1 (g), and 0 (h) repressively modified nucleosomes. After ACC and HS, a considerable fraction of cells are suggested to stay at states with 1 or 2 actively modified nucleosomes. These results indicate the importance in the memory effect of having multiple modifiable sites.

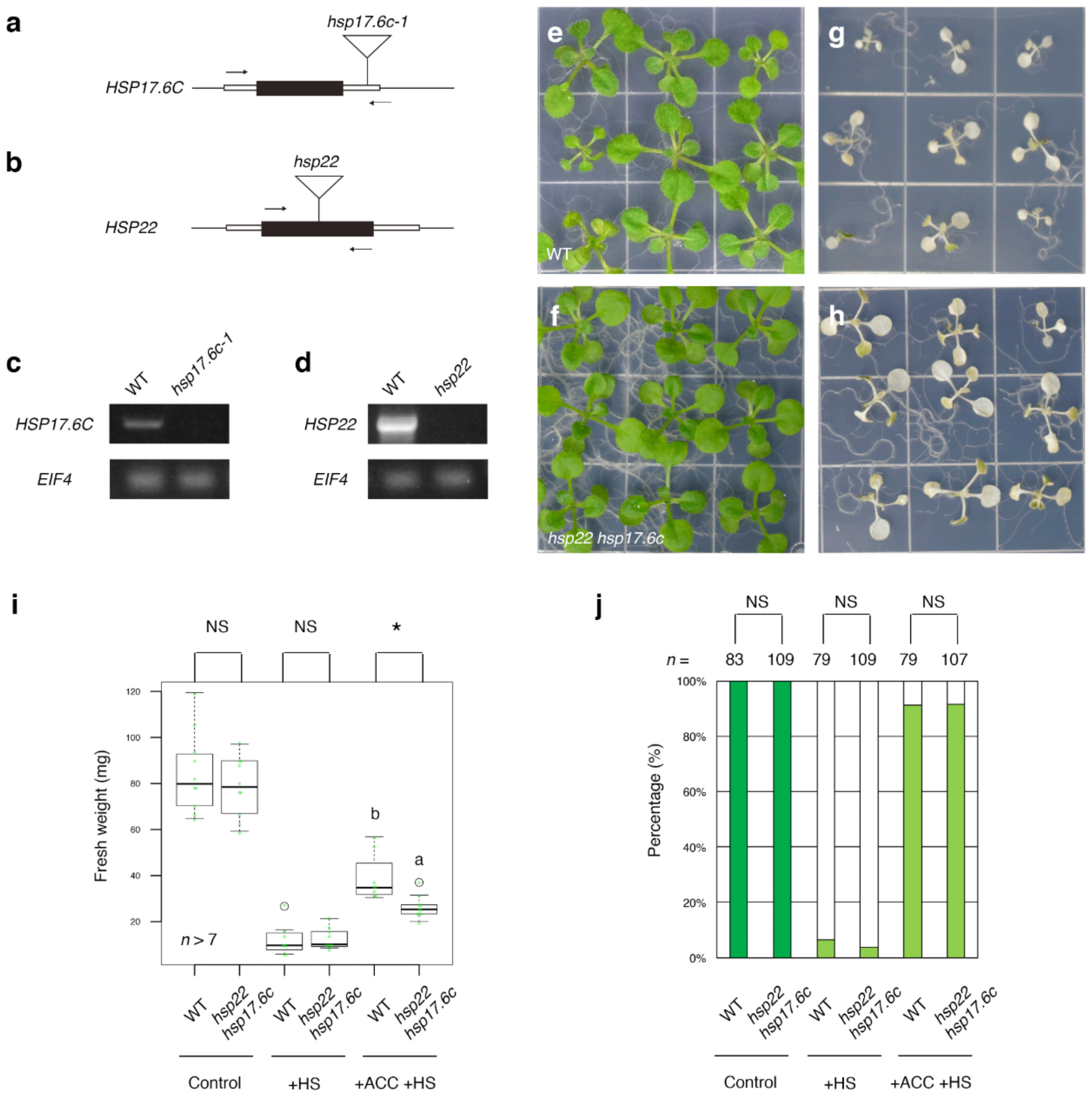


**Extended Data Fig. 13 The *hsp22 hsp17.6c* double mutant showed decreased heat acclimation capacity.**

**a** and **b**, Schematic structure of *HSP17.6C* (a) and *HSP22* (b) genes and T-DNA insertions. Arrows indicate gene-specific primers used for genotyping. **c**, Expression of *HSP17.6C* (above) and *EIF4A1* (below) in wild-type and *hsp17.6c-1* seedlings. *EIF4A1* was used for a loading control. **d**, Expression of *HSP22* (above) and *EIF4A1* (below) in wild-type and previously identified *hsp22* seedlings[15]. *EIF4A1* was used for a loading control. **e** and **f**, Wild-type (e) and *hsp22 hsp17.6c* (f) seedlings grown under control condition. **g** and **h**, Wild type (g) and *hsp22 hsp17.6c* (h) grown under +HS condition. **i**, Quantification of seedling fresh weights. Asterisk indicates significant differences based on one-way ANOVA test. * $p < 0.05$. NS, nonsignificant. Different letters indicate significant differences, while the same letters indicate non-significant differences based on post-hoc Tukey's HSD test. $p < 0.05$. **j**, Quantification of survival rate. Ten-day-old seedlings grown under three different temperature conditions were categorized into three groups based on phenotypic severity: green, normal growth; light green, partially damaged; white, perished. Significance was determined by $\chi^2$ test and the post-hoc test that followed. NS, nonsignificant. $n > 78$. Although fresh weight in acclimaized *hsp22 hsp17.6c* double mutants was significantly lighter than that in acclimaized wild type after heat shock, no difference in survival rate was observed. The lack of difference in survival rate of *hsp22 hsp17.6c* double mutants, while there is a difference in *jmjq* mutants, suggests that other differentially expressed genes, such as *HSP21* may also contribute to phenotypic consequence of *jmjq* mutants for heat acclimation.

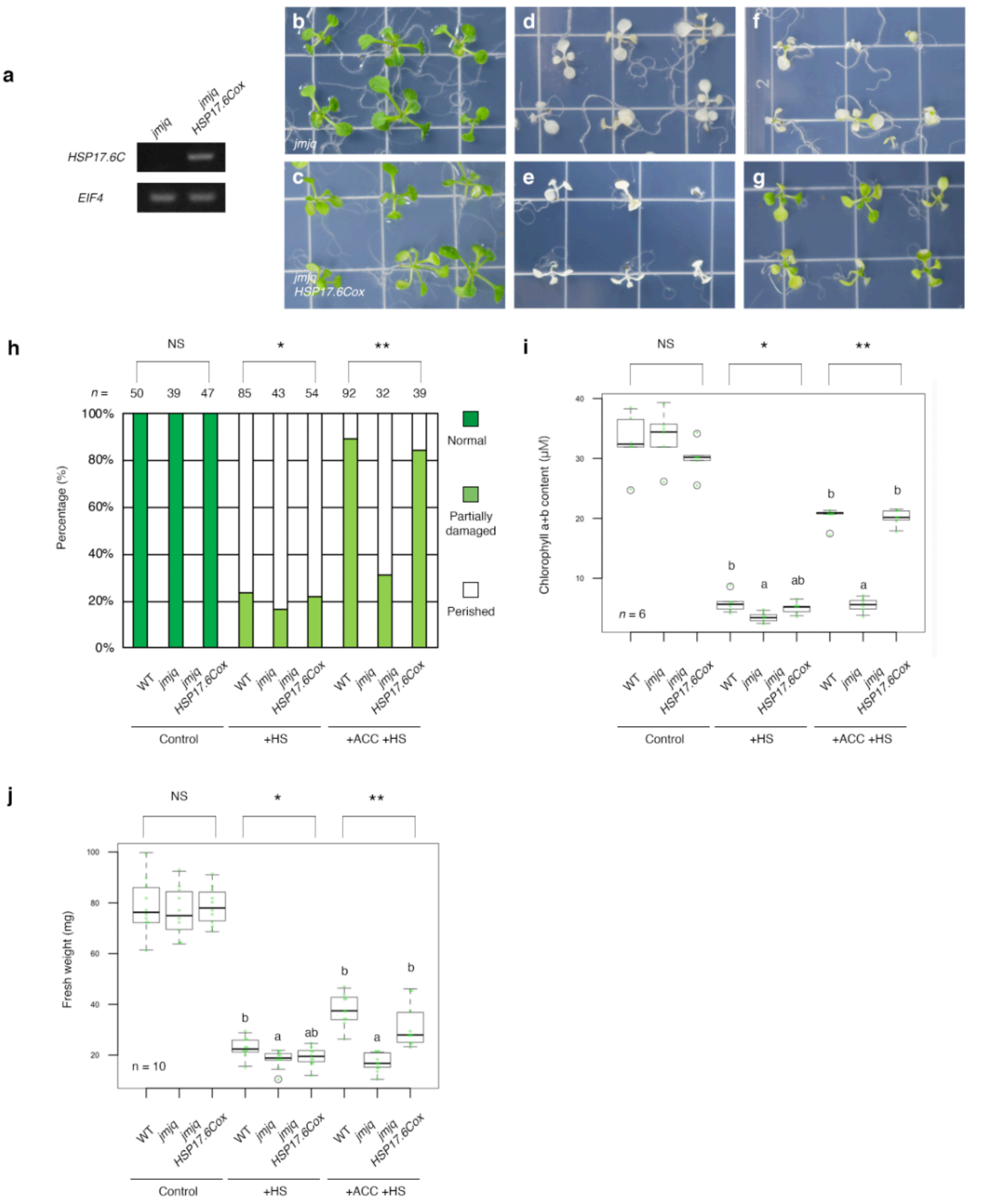


**Extended Data Fig. 14 Ectopic expression of *HSP17.6C* rescued the heat acclimation phenotype in *jmjq* mutants.**

**a**, Expression of *HSP17.6C* (above) and *EIF4A1* (below) in *jmjq* and *jmjq 35S::HSP17.6C* seedlings. *EIF4A1* was used for a loading control. **b** and **c**, *jmjq* (b) and *jmjq 35S::HSP17.6C* (c) grown under control condition. **d** and **e**, Wild type (d) and *jmjq* (e) grown under +HS condition. **f** and **g**, Wild type (f) and *jmjq*

(g) grown under +ACC +HS condition. **h**, Quantification of survival rate. Ten-day-old seedlings grown under three different temperature conditions were categorized into three groups based on phenotypic severity: green, normal growth; light green, partially damaged; white, perished. Significance was determined by $\chi^2$ test and the post-hoc test that followed. $n > 31$. **i**, Quantification of chlorophyll contents. Light green jitter dots and white circles represent the chlorophyll content from each sample and statistical outliers, respectively. Asterisks indicate significant differences based on one-way ANOVA test. *$p < 1.0 \times 10^{-2}$, **$p < 1.0 \times 10^{-3}$. Different letters indicate significant differences, while the same letters indicate non-significant differences based on post-hoc Tukey's HSD test. $p < 0.05$. NS, nonsignificant. **j**, Quantification of seedling fresh weights. Asterisks indicate significant differences based on one-way ANOVA test. *$p < 5.0 \times 10^{-2}$, **$p < 1.0 \times 10^{-4}$. Different letters indicate significant differences, while the same letters indicate non-significant differences based on post-hoc Tukey's HSD test. $p < 0.05$. NS, nonsignificant.

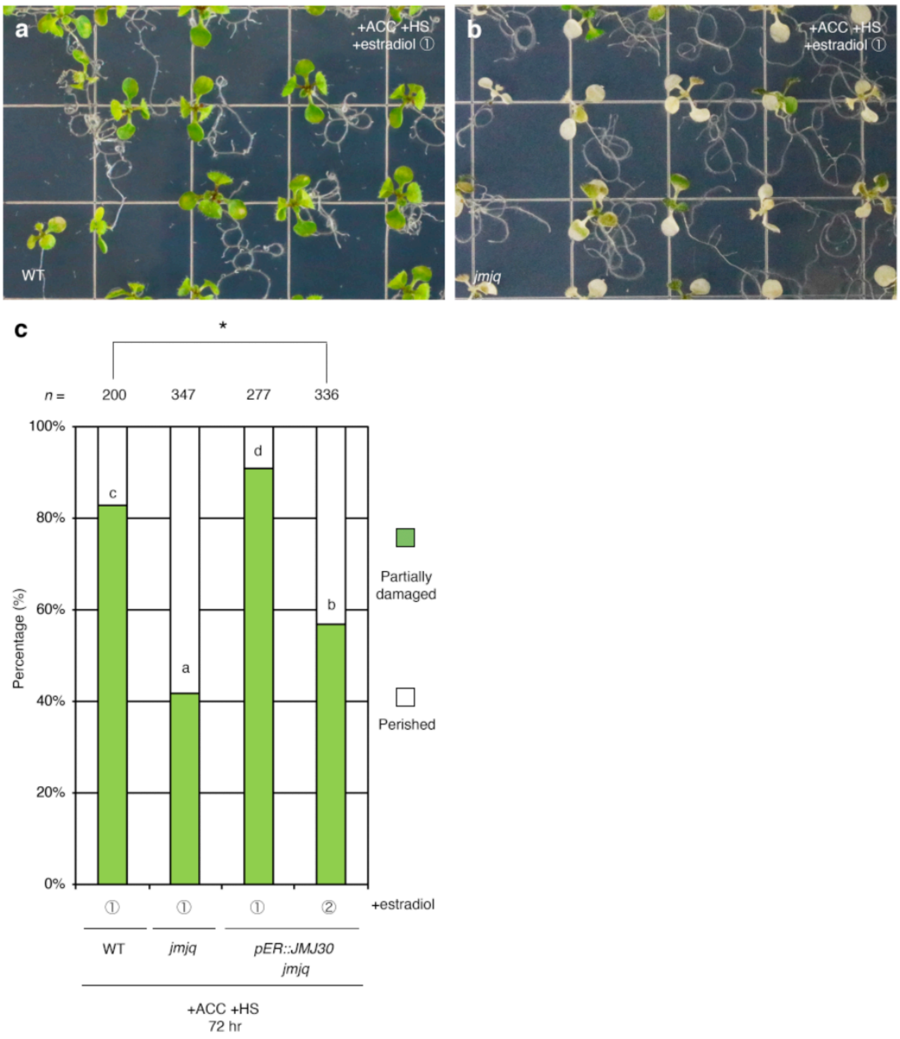


**Extended Data Fig. 15 Induction of *JMJ30* in *jmjq* mutants prior to acclimation rescues the mutant phenotype. a** and **b**, Images of wild type (a) and the *jmjq* mutant (b) with β-estradiol application before acclimation (left) and before heat shock (right). Plants were grown under +ACC +HS condition. **c**, Quantification of survival rate shown in Fig. 4b and Extended Data Fig. 15a and b. Ten-day-old seedlings grown under +ACC +HS condition with β-estradiol application before acclimation and before heat shock were categorized into three groups based on phenotypic severity: green, normal growth; light green, partially damaged; white, perished. Significance was determined by $\chi^2$ test and the post-hoc test that followed. *n* > 199.

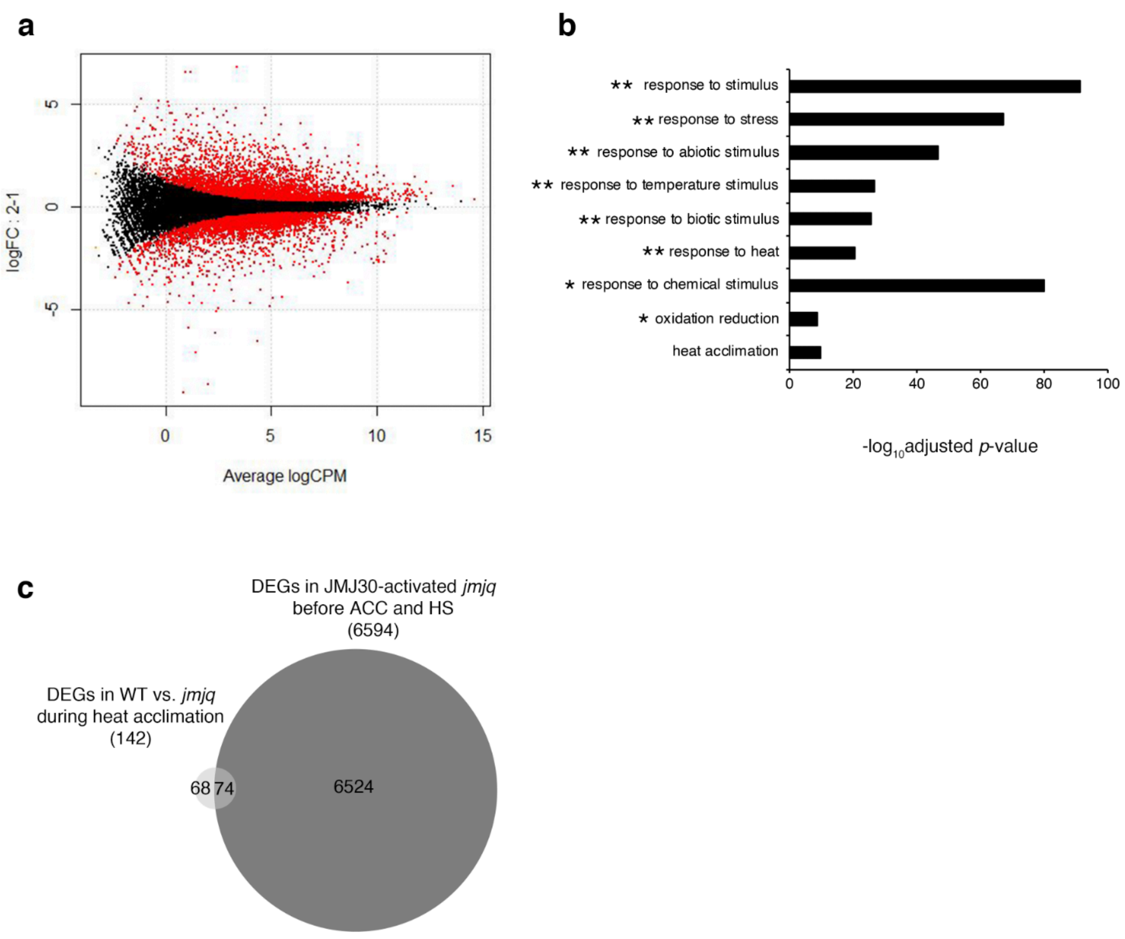


**Extended Data Fig. 16 Induction of *JMJ30* in *jmjq* mutants prior to acclimation triggers heat-acclimation-related gene expression. a**, The MA plots represent each gene with a dot. The *x* axis is average log CPM over all genes; the *y* axis is the $\log_2$ fold change of normalized count between *pER8::JMJ30* transgenic plants in the *jmjq* mutant background with β-estradiol application before acclimation and before heat shock. Genes with FDR <0.05 are shown in red. 6594 genes were differentially expressed when *JMJ30* was misexpressed before acclimation or before heat shock. **b**, Gene ontology (GO) term enrichment analysis of 6594 genes. Selected GO terms determined by their $-\log_{10}$-adjusted *p*-values are shown. All enriched GO terms are shown in Data S8. **, GO terms seen in Fig. 2b; *, GO terms similar to those observed in Fig. 2B. Similar pathways were affected in *jmjq* mutants and JMJ30-induced plants prior to acclimation. **c**, Venn diagram showing the overlap between differentially expressed genes in wild type and *jmjq* mutants, and differentially expressed genes in *pER8::JMJ30* transgenic plants in the *jmjq* mutant background with β-estradiol application before acclimation and before heat shock. This overlap was significantly larger than expected by chance ($p = 3.0 \times 10^{-18}$). The 74 overlapping genes included *HSP17.6C*, *HSP21*, and *HSP22* as well as *HSP23.6-MITO*, *HSP17.6 II*, *HSP70B*, and *HSP70-T2*.

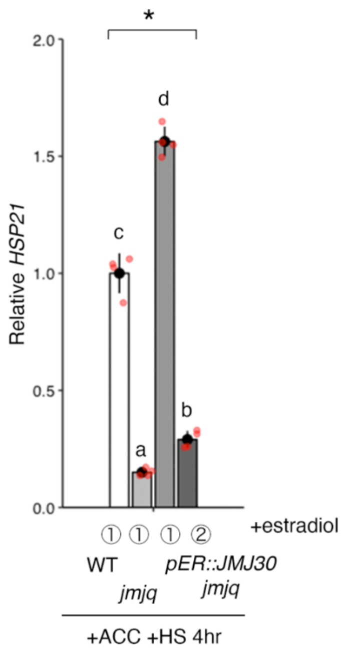


**Extended Data Fig. 17 *HSP21* expression in *JMJ30*-induced *jmjq* mutants prior to acclimation.** Gene expression levels of *HSP21* in the wild type, *jmjq* mutants, and *pER8::JMJ30* transgenic plants in the *jmjq* mutant background with β-estradiol application before acclimation and before heat shock. Asterisk indicates significant differences based on one-way ANOVA test. Different letters indicate significant differences, while the same letters indicate non-significant differences based on post-hoc Tukey's HSD test. $p < 0.05$.

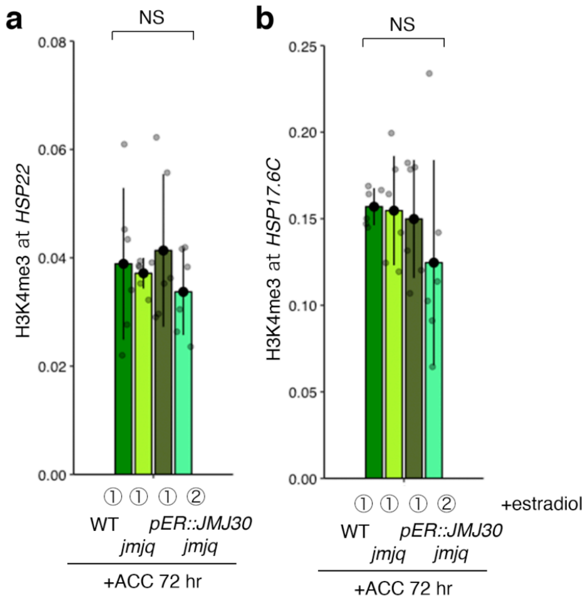


**Extended Data Fig. 18 Histone modifications at the *HSP22*, and *HSP17.6C* loci in *JMJ30*-induced *jmjq* mutants prior to acclimation. a** and **b**, H3K4me3 levels at *HSP22* (a) and *HSP17.6C* (b) in the wild type, *jmjq* mutants, and *pER8::JMJ30* transgenic plants in the *jmjq* mutant background with β-estradiol application before acclimation and before heat shock. NS, nonsignificant based on one-way ANOVA test.

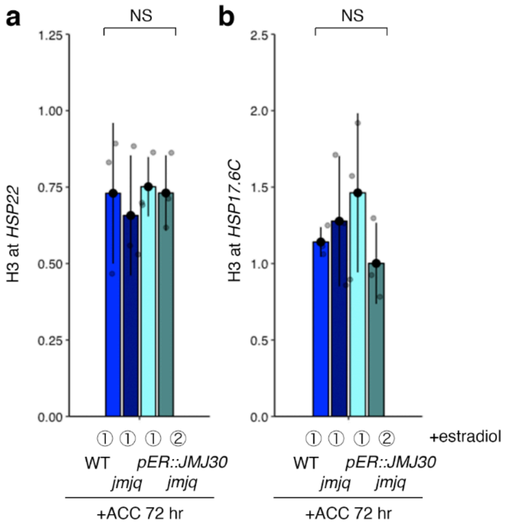


**Extended Data Fig. 19 Histone H3 levels at the *HSP22*, and *HSP17.6C* loci in *JMJ30*-induced *jmjq* mutants prior to acclimation. a** and **b**, H3 levels at *HSP22* (a), and *HSP17.6C* (b) in the wild-type, *jmjq* mutants, and *pER8::JMJ30* transgenic plants in the *jmjq* mutant background with β-estradiol application before acclimation and before heat shock. NS, nonsignificant based on one-way ANOVA test. Taken together with the H3K27me3 and H3K4me3 results shown in Fig. 4, these data imply that it is histone modifications, rather than the locations of histones (or nucleosomes) along the DNA, that are changed due to acclimation.

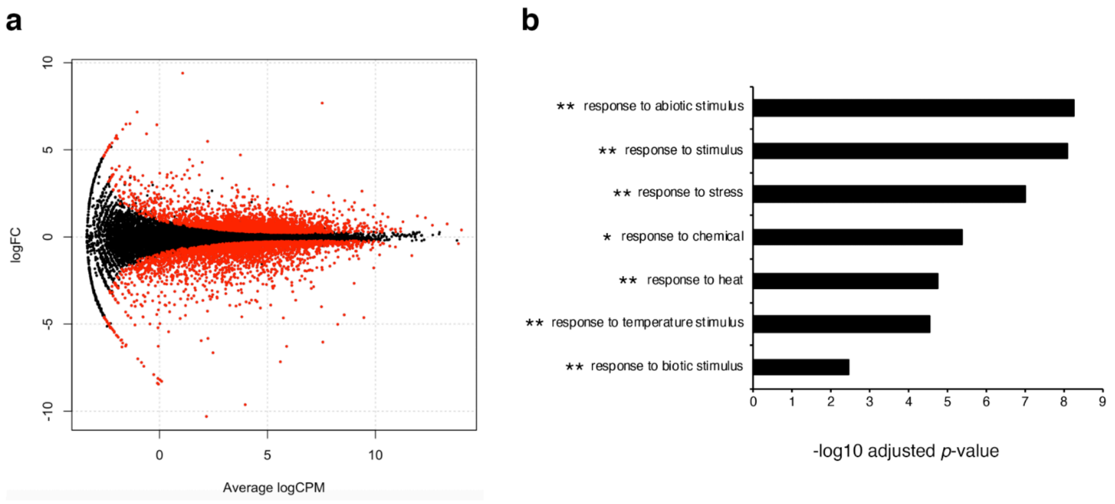


**Extended Data Fig. 20 Gene expression in wild type and *jmjq* mutants under the Nara condition. a**, The MA plots represent each gene with a dot. The *x* axis is average log CPM over all genes; the *y* axis is the $\log_2$ fold change of normalized count between wild type and *jmjq* mutants under the Nara condition. Genes with FDR <0.05 are shown in red. 5947 genes were differentially expressed. **b**, Gene ontology (GO) term enrichment analysis of 46 genes. Selected GO terms determined by their $-\log_{10}$-adjusted *p*-values are shown. All enriched GO terms are shown in Supplementary Table 12. **, GO terms seen in Fig. 2b; *, GO terms similar to those observed in Fig. 2b. Similar pathways were affected in *jmjq* mutants and JMJ30-induced plants prior to acclimation.

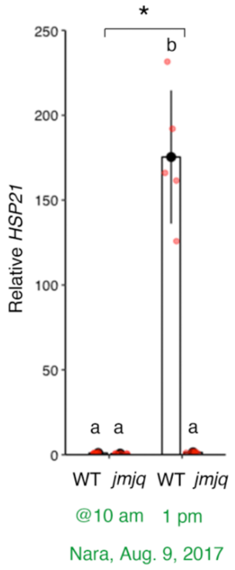


**Extended Data Fig. 21 *HSP21* expression in wild type and *jmjq* mutants under the Nara condition.** Gene expression levels of *HSP21* in the wild-type and *jmjq* mutants under the Nara condition. Asterisk indicates significant differences based on one-way ANOVA test. Different letters indicate significant differences, while the same letters indicate non-significant differences based on post-hoc Tukey's HSD test.

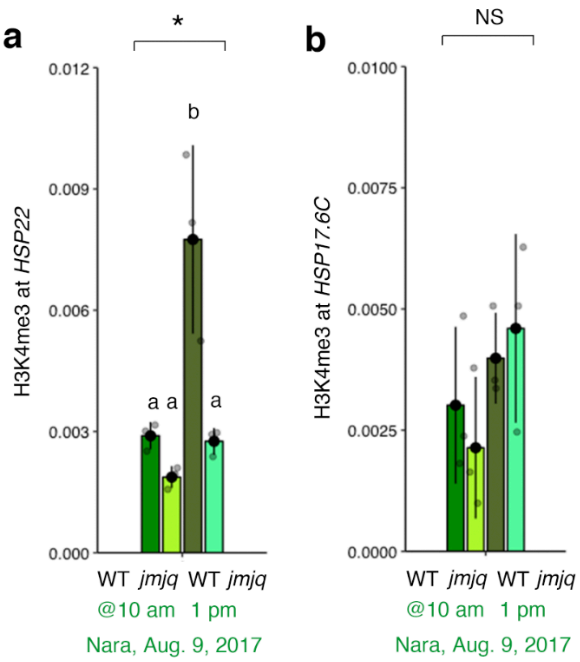


**Extended Data Fig. 22 Histone modifications at the *HSP22*, and *HSP17.6C* loci under the Nara condition. a** and **b**, H3K4me3 levels at *HSP22* (a), and *HSP17.6C* (b) in the wild type and *jmjq* mutants under the Nara condition. Asterisk indicates significant differences based on one-way ANOVA test. Different letters indicate significant differences, while the same letters indicate non-significant differences based on post-hoc Tukey's HSD test. $p < 0.05$. NS, nonsignificant. The difference in H3K4me3 enrichment between *HSP22* and *HSP17.6C* in wild type at 1 pm under the Nara condition suggests that conditions are important for the regulation of *HSP22* and *HSP17.6C* by histone demethylases.

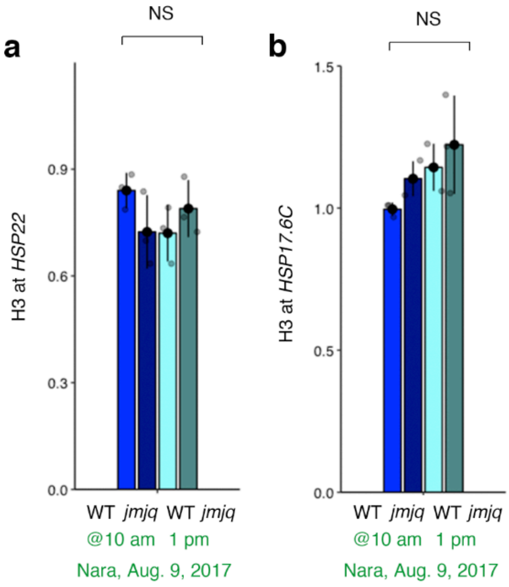


**Extended Data Fig. 23 Histone H3 levels at the *HSP22*, and *HSP17.6C* loci under the Nara condition. a** and **b**, H3 levels at *HSP22* (a), and *HSP17.6C* (b) in the wild type and *jmjq* mutants under the Nara condition. NS, nonsignificant based on one-way ANOVA test. Taken together with the H3K27me3 results shown in Fig. 4, these data imply that it is histone modifications, rather than the locations of histones (or nucleosomes) along the DNA, that are changed due to acclimation.

Supplementary Note and Table 1 and 2 for

### Removal of repressive histone marks creates epigenetic memory of recurring heat in *Arabidopsis*

**Authors:** Nobutoshi Yamaguchi[1,2*], Satoshi Matsubara[1], Kaori Yoshimizu[1], Motohide Seki[3], Kouta Hamada[4], Mari Kamitani[5], Yuko Kurita[5], Soichi Inagaki[2,6], Takamasa Suzuki[7], Eng-Seng Gan[8], Taiko To[6], Tetsuji Kakutani[6,9,10], Atsushi J. Nagano[5,9], Akiko Satake[4], Toshiro Ito[1*]

**Definition of transition rates between three states of histone modification**

We developed a mathematical model describing state transitions of histone modification at the cell-population level to predict the level of *HSP* expression under changing temperature.

We focus on a locus with $N$ units of nucleosomes. Each nucleosome is in one of the following three states, actively modified (A), unmodified (U), and repressively modified (R) (Fig. 3g), as assumed in a previous study[19,20]. In an actively transcribed *HSP* chromatin, H3K4me3 modifications are enriched. The unmodified state changes to the repressively modified state by the addition of repressive histone marks. H3K27me3 marks are associated with the repressed transcriptional state of genes. Let $i$ and $j$ be the numbers of actively- and repressively-modified nucleosomes, respectively ($0 \leq i,j \leq N$ and $i + j \leq N$). The number of unmodified nucleosomes is calculated as $N - i - j$, and thus every possible state of the locus can be described with a pair of integers $(i,j)$. Transcription of the locus occurs only when all $N$ nucleosomes are in state A (i.e., when $(i,j) = (N,0)$).

State transition of each nucleosome is modeled based on a previous study[20]. Consider a sufficiently short period of time during which at most one of the $N$ units in each locus can change its state. Let a transition rate at which a nucleosome in the state X changes to the state Y as $r_{X\to Y}$, where $(X,Y) \in \{(\mathrm{U},\mathrm{A}),(\mathrm{R},\mathrm{U}),(\mathrm{U},\mathrm{R}),(\mathrm{A},\mathrm{U})\}$. Due to positive feedback mechanisms involved in the recruitment of histone modifying complexes, all-A and all-R modification states can be stable at the same time; that is, the system can be 'bistable' [19,20]. This positive feedback is incorporated in the model by assuming transition rates as a function of the number of nucleosomes in the state A or R. To be specific, transition rate from the state A or U to the state A increases when the number of A within the focal locus ($i$) is greater. It follows that $r_{\mathrm{U}\to\mathrm{A}}$ and $r_{\mathrm{R}\to\mathrm{U}}$ are increasing functions of $i$ as follows:

$$r_{\mathrm{U}\to\mathrm{A}}(i;T) = v_{\mathrm{U}\to\mathrm{A}} \cdot P(T) \cdot (1 + \beta_{\mathrm{U}\to\mathrm{A}} \cdot i) \qquad \text{(Eq. 1a)}$$

$$r_{\mathrm{R}\to\mathrm{U}}(i;T) = v_{\mathrm{R}\to\mathrm{U}} \cdot P(T) \cdot (1 + \beta_{\mathrm{R}\to\mathrm{U}} \cdot i), \qquad \text{(Eq. 1b)}$$

where $v_{X\to Y}$ is a transition rate when the number of A equals zero under ambient temperature of 22˚C. The function $P(T)$ represents the dependence of transition rates on temperature ($T$). High temperature is expected to induce expression of *HSP* genes, and we define $P(T)$ as follows:

$$\begin{cases} P(22) = 1 & \text{under ambient temperature of } 22^\circ\text{C} \\ P(37) = P_{\mathrm{ACC}} & \text{during acclimation under } 37^\circ\text{C} \\ P(44) = P_{\mathrm{HS}} & \text{during heat shock under } 44^\circ\text{C} \end{cases}.$$

Coefficients $\beta_{X\to Y}$ represent the strength of positive feedback. This linear formulation is not a strong assumption, as we obtain similar results as long as the feedback term is a monotonically increasing function of $i$.

Similarly, $r_{\mathrm{U\to R}}$ and $r_{\mathrm{A\to U}}$ are given as increasing functions of the number of R ($j$) as follows:

$$r_{\mathrm{U\to R}}(j) = v_{\mathrm{U\to R}} \cdot (1 + \beta_{\mathrm{U\to R}} \cdot j) \qquad \text{(Eq. 2a)}$$

$$r_{\mathrm{A\to U}}(j) = v_{\mathrm{A\to U}} \cdot (1 + \beta_{\mathrm{A\to U}} \cdot j), \qquad \text{(Eq. 2b)}$$

where $v_{X\to Y}$ and $\beta_{X\to Y}$ are defined similarly as in Eq. (1).

**Dynamics of histone modification at the cell-population level**

Using the transition rate functions defined in Eqs. (1) and (2), we describe the dynamics describing state change of histone modification at the cell-population level by assuming no interaction between cells. Let $s_{i,j}(t)$ be the proportion of cells in the state $(i,j)$ ($\sum_{i=0}^{N}\sum_{j=0}^{N-i} s_{i,j}(t) = 1$). Some fraction of cells in the state $(i,j)$ will transition to the state $(i+1,j)$ by changing the state of one nucleosome from U to A at a rate $r_{\mathrm{U\to A}}(i;T)$. At the same time, some of cells in the state $(i+1,j)$ will shift into the state $(i,j)$ by reducing the number of A by one at a rate $r_{\mathrm{A\to U}}(i+1)$. The above outflow and inflow between the states $(i,j)$ and $(i+1,j)$ is summarized in a mathematical form as $I^+$:

$$I^+ = -r_{\mathrm{U\to A}}(i;T) \cdot s_{i,j}(t) + r_{\mathrm{A\to U}}(i+1) \cdot s_{i+1,j}(t). \qquad \text{(Eq. 3)}$$

Likewise, the subpopulation of cells in the state $(i,j)$ can have outflow toward and inflow from another neighboring state $(i,j+1)$, which is denoted by $J^+$:

$$J^+ = -r_{\mathrm{U\to R}}(j) \cdot s_{i,j}(t) + r_{\mathrm{R\to U}}(j+1;T) \cdot s_{i,j+1}(t). \qquad \text{(Eq. 4)}$$

Note that Eqs. (3) and (4) presume that the state $(i,j)$ involves at least one unmodified unit (i.e., $N-i-j > 0$ or $i+j < N$). Similarly, if cells are in states with at least one nucleosome in state A (i.e., $i \geq 1$) there are flows between the state $(i,j)$ and its neighboring state $(i-1,j)$:

$$I^{-} = -r_{\mathrm{A}\to\mathrm{U}}(i;T)\cdot s_{i,j}(t) + r_{\mathrm{U}\to\mathrm{A}}(i-1;T)\ \cdot s_{i-1,j}(t) \qquad \text{(Eq. 5)}$$

In the end, flows exist between the state with at least one nucleosome in state R ($j \geq 1$) and its neighboring state $(i, j-1)$:

$$J^{-} = -r_{\mathrm{R}\to\mathrm{U}}(j;T)\cdot s_{i,j}(t) + r_{\mathrm{U}\to\mathrm{R}}(j-1)\ \cdot s_{i,j-1}(t) \qquad \text{(Eq. 6)}$$

Using Kronecker's $\delta_{x,y}$ ($\delta_{x,y} = 1$ if $x = y$ and $\delta_{x,y} = 0$ otherwise), we have

$$\frac{\mathrm{d}}{\mathrm{d}t} s_{i,j} = \left(1-\delta_{i+j,N}\right)(I^{+} + J^{+}) + \left(1-\delta_{i,0}\right)I^{-} + \left(1-\delta_{j,0}\right)J^{-}. \qquad \text{(Eq. 7)}$$

It can be mathematically shown that the above system has a unique and globally stable equilibrium under a constant temperature. We define the equilibrium state at $T = 22$ (°C) as $\left(\hat{s}_{0,0}, \hat{s}_{0,1}, \dots, \hat{s}_{N,0}\right)$.

Because we assume that *HSP* genes are expressed only when all the $N$ nucleosomes are in state A, expression levels of *HSP* genes at time $t$, denoted as $m(t)$, are in proportion to the fraction of cells in the state (*N*, 0), $s_{N,0}(t)$, as follows:

$$m(t) = q \cdot s_{N,0}(t), \qquad \text{(Eq. 8)}$$

where $q$ is a positive constant. In the following analyses, we substitute $q = 1/\hat{s}_{N,0}$ so that $m(0) = 1$. Although we do not observe cell death, a majority of cells are damaged after heat shock treatment (Extended Data Fig. 6). Thus, we assume that the fraction $h$ of cells stop transcription of *HSP* genes right after the heat shock at $t = t_{\mathrm{HS}}$ and gradually recover their transcription activity at a constant rate $r$:

$$m(t) = \begin{cases} q \cdot s_{N,0}(t) & \text{for } t \leq t_{\mathrm{HS}} \\ q \cdot \left(1 - h \cdot e^{-r\cdot(t - t_{\mathrm{HS}})}\right) \cdot s_{N,0}(t) & \text{for } t > t_{\mathrm{HS}} \end{cases}. \qquad \text{(Eq. 9)}$$

Because the ORF length of *HSP22* is 588 bp, we choose $N = 3$ for the following analyses. In our formalization, we assume that the order of histone modification at different nucleosomes is not random, but there is a specific order (e.g. a nucleosome closest to the transcription start site is the first to be repressively modified.). For parameter fitting using wild type, the sum of squared errors between log-transformed experimental and simulation data was considered as a cost function to be minimized. The present model has as many as 12 free parameters, in which case it is generally difficult to specify the single optimal solution. Therefore, we used 144 different sets of initial parameter values that were randomly chosen from fixed ranges for each parameter (Table 1) and obtained the best fit set of parameters (Table 2). For parameter fitting, time unit is set to an hour. We also performed parameter fitting using experimental data from *jmjq* mutant. We chose the wild-type best-fit parameters as initial values for this analysis on the basis that a mutant would not be drastically different from wild type. Comparison of best fit parameters between wild type and *jmjq* mutant showed that the *jmjq* mutant has notably greater values for the strength of positive feedback for the transition from A to U ($\beta_{\mathrm{A}\to\mathrm{U}}$) and temperature dependence during heat shock ($P_{\mathrm{HS}}$). On the contrary, the *jmjq* mutant has smaller values for recovery rate of transcription activity after heat damage ($r$) and the basic transition rate from R to U ($v_{\mathrm{R}\to\mathrm{U}}$). The greater positive feedback effect ($\beta_{\mathrm{A}\to\mathrm{U}}$) and the smaller basic transition rate from R to U corresponds to the feature that the *jmjq* mutant lacks the demethylation activity for the repressive mark H3K27me3.

The model can predict dynamics of histone modification states that have not measured in the experiments (Extended Data Fig. 12). The results also revealed the importance to the memory effect of having multiple modifiable sites.

**Predicting *HSP* expression profiles in natural temperature conditions**

We also performed simulations with temperature profiles obtained from several fields. Based on the suggestion from the above parameter fittings that $P(T)$ is a nonlinear, monotonically increasing function of temperature, we assume that $P(T)$ takes the form of exponential function:

$$P(T) = k_1 + k_2 \cdot e^{k_3 \cdot T}, \qquad \text{(Eq. 10)}$$

where $k_1$, $k_2$, and $k_3$ are constant parameters. Substituting the definition $P(22) = 1$ and the best-fit parameter values, $P(37) = P_{\mathrm{ACC}}$ and $P(44) = P_{\mathrm{HS}}$, we obtained estimated values for those coefficients ($k_1 = 3.61 \times 10^{-6}$, $k_2 = 0.355$, and $k_3 = 0.911$). We interpolated hourly temperature data at Nara and substituted it into the present model to obtain Fig. 4h.

We used *Mathematica* 11.3.0.0 (Wolfram Research Inc.) for numerical simulations of the ordinary differential equations (Adams' method), parameter fitting (the interior point method), and interpolation (to sixth order polynomial).

**Table 1. Ranges of random values from which initial parameter values were chosen.**

| | Lower limit | Upper limit |
|---|---|---|
| $v_{X \to Y}$ | 0.1 | 2 |
| $\beta_{\mathrm{X \to Y}}$ | 0.1 | 7 |
| $P_{\mathrm{ACC}}, P_{\mathrm{HS}}$ | 1 | 100 |
| $h, r$ | 0.1 | 0.9 |

**Table 2. Best-fit parameter values.**

| | WT parameters | *jmjq* parameters | (*jmjq* − WT) / WT |
|---|---|---|---|
| $v_{\mathrm{U\to A}}$ | 1.6220 | 1.6156 | − |
| $v_{\mathrm{R\to U}}$ | 0.063225 | 0.050555 | $-20\%$ |
| $v_{\mathrm{U\to R}}$ | 4.8494 | 4.8672 | − |
| $v_{\mathrm{A\to U}}$ | 0.21465 | 0.20279 | $-5\%$ |
| $\beta_{\mathrm{U\to A}}$ | 1.2618 | 1.2673 | $+9\%$ |
| $\beta_{\mathrm{R\to U}}$ | 0.49607 | 0.54300 | − |
| $\beta_{\mathrm{U\to R}}$ | 4.6667 | 4.6891 | − |
| $\beta_{\mathrm{A\to U}}$ | 8.2086 | 10.426 | $+27\%$ |
| $P_{\mathrm{ACC}}$ | 19.134 | 19.132 | − |
| $P_{\mathrm{HS}}$ | 219.44 | 263.40 | $+20\%$ |
| $h$ | 1 | 1 | − |
| $r$ | 0.0099957 | 0.0077896 | $-22\%$ |
| Value of cost function | 10.70 | 21.74 | − |

Note: Values within $\pm1\%$ are not shown in the far-right column.
